# Climate-linked divergence in tree flowering and fruiting in an Eastern Himalayan tropical forest

**DOI:** 10.64898/2026.02.26.708186

**Authors:** Soumya Banerjee, Aparajita Datta

## Abstract

**Premise:** Long-term phenological patterns are increasingly being examined from the perspective of climate change and its potential effects. Climatic effects on plant phenology could involve the direct responses to changes in temperature, precipitation and solar irradiance, or could be mediated by these variables through exogenous teleconnections such as the El Niño Southern Oscillation (ENSO). The effects of climatic fluctuations on inter-annual variation in tropical phenology remain understudied.

**Methods:** We examined long-term patterns of tree flowering and fruiting intensity in a tropical forest site in the Eastern Himalayas between 2011 and 2024. Species-specific patterns were examined for 36 species. Long-term patterns were quantified using Generalized Additive Models, and splines were visualized to infer trends. Through Generalized Linear Mixed Models, we determined if there was a lagged phenological response to ENSO and temperature, precipitation and solar irradiance, and whether ENSO effects were being mediated through the latter group of variables or plant traits.

**Results:** Between 2011 and 2019, trends in flowering and fruiting were significant for 17 and 23 species respectively. Flowering increased for 7 species, while fruiting declined for 8 species. Flowering peaked during El Niño, but this association did not appear to be mediated through climate variables, whereas fruiting showed a three-month positive lagged response to solar irradiance, independent of ENSO. The peak season of reproduction was the only trait determining species-specific responses to climate variables.

**Conclusion:** Our study highlights nonlinearity in long-term patterns of reproductive phenology, and the importance of solar irradiance in determining inter-annual fruit production.

## INTRODUCTION

Climatic variables such as temperature, precipitation, solar irradiance and daylength are the main proximate cues of plant vegetative and reproductive phenology (van Schaik et al. 1993, Chapman et al. 1999, Borchert et al. 2002, Wright & Calderón 2018). In the tropics, owing to fewer temperature-related constraints on plant physiology, daylength and solar irradiance are the principal proximate cues of leafing, flowering and fruiting in habitats with adequate year-round soil moisture (Elliot et al. 2006, Zimmerman et al. 2007, Datta et al. 2025). The absence of a clearly defined growing season in the tropics results in a much higher diversity in phenological patterns (Sakai 2001). Thus, tropical species may show distinct responses in phenological patterns to anthropogenic climate change compared to those at higher latitudes. For instance, the widely reported temperate-zone advancement of spring phenology (Parmesan & Yohe 2003, Primack et al. 2004, Karlsen et al. 2009, Primack et al. 2009) may not be mirrored in the tropics (e.g. Mendoza et al. 2018). However, tropical species are broadly expected to be more vulnerable to climate change owing to their adaptation to narrower thermal niches (Deutsch et al. 2008, Tewksbury et al. 2008, Linck et al. 2021).

Phenological responses to climate change are also likely to vary among species across taxonomic groups, and this could be mediated by morphological and phenological traits (Fei et al. 2017, Pacifici et al. 2017, Germain et al. 2023). For instance, species flowering earlier in the season are often known to advance their flowering dates faster in response to climate change (Fitter & Fitter 2002, Miller-Rushing & Inouye 2009).

The majority of studies on the effects of climate change on plant phenology have examined shifts in the timing of phenological events, such as budburst and flowering (e.g. Miller-Rushing et al. 2008, Büntgen et al. 2022, Rana et al. 2025). Long-term changes in phenological intensity, through metrics such as the abundance of flowers or fruit (e.g. Chapman et al. 2005, Dunham et al. 2018), have been relatively understudied, even though inter-annual variation in reproductive intensity may be more significant than changes in phenological timing in response to climate change (e.g. Mendoza et al. 2018).

Long-term trends in flowering intensity and abundance have been particularly understudied in the tropics. Increases in flowering have been reported from tropical forest sites in Barro Colorado, Panama (Wright & Calderón 2006, Pau et al. 2018) and among lianas in Australia (Vogado et al. 2022). The few studies on long-term fruiting patterns in the tropics have produced equivocal results. In Uganda (Chapman et al. 2005, Potts et al. 2020), and Cote d’Ivoire (Polansky & Boesch 2013), an increase in fruiting was observed. On the other hand, in Gabon, fruit availability declined (Bush et al. 2020). In French Guiana, there was no discernible trend in fruiting, although there was substantial inter-annual variation in fruit productivity (Mendoza et al. 2018). Likewise, long-term patterns from Panama did not show an increase in fruiting, even though there was an increase in flowering (Wright & Calderón 2006). The overall global trend appears to be one of declining fruit availability over time (Hacket-Pain et al. 2024). Trends in reproductive intensity in the tropics are typically non-monotonic and can be complex (e.g Polansky & Robbins 2013, Potts et al. 2020).

The mechanistic pathways by which climate patterns affect inter-annual variability in tree phenology have been understudied. Warmer and wetter years appeared to be associated with an increase in fruiting in Uganda (Chapman et al. 2018), but Polansky & Boesch (2013) did not find a relationship between inter-annual variation in rainfall and fruiting. In sites where phenology is strongly seasonal, the same climatic cues can be expected to determine both seasonal and inter-annual variability (Wright & Calderón 2006). However, climatic extremes could disrupt this relationship. For instance, periodic severe droughts and the resultant water stress could make inter-annual phenology more dependent on rainfall (Pau et al. 2013), even if the predominant cues of phenology in seasonal tropical forests are daylength and solar irradiance (Elliot et al. 2006, Zimmerman et al. 2007, Datta et al. 2025). In addition to inter-annual climatic variation, an increase in CO_2_ has been found to directly increase flower abundance in Barro Colorado, Panama (Pau et al. 2018) and more generally, an increase in energetic allocation to reproduction in plants (Jablonski et al. 2002).

One potential mechanistic pathway determining inter-annual variation in tree phenology involves the effect of climatic teleconnections such as the El Niño Southern Oscillation (ENSO) on temperature and precipitation, which in turn, could affect phenological patterns. ENSO is a climatic phenomenon of complex periodicity (Kestin et al. 1998, Ren & Wang 2023) associated with anomalies in sea surface temperature in the Eastern Pacific that has significant effects on atmospheric climatic conditions across the tropics (Horel & Wallace 1981, Yeh et al. 2018, Cai et al. 2020).

In South Asia, droughts resulting from reduced rainfall during the southwest monsoon have historically been associated with positive ENSO anomalies (or El Niño episodes, while anomalously reduced sea surface temperatures are referred to as the “La Niña”) (Mooley & Parthasarathy 1983, Kumar et al. 1995) through changes in the Walker circulation (Soman & Slingo 1997, Dai & Wigley 2000). However, this relationship has been disputed by recent research. The southwest monsoon rainfall may be more sensitive to sea surface temperature anomalies in the Central Pacific, than those in the Eastern Pacific (Kumar et al. 2006). In addition, the strength of the ENSO-SW monsoon coupling tends to vary over time (Kumar et al. 1999, Athira et al. 2023). Teleconnections in the Indian Ocean region such as the Indian Ocean Dipole (IOD) can also mediate the degree of ENSO forcing (Ashok et al. 2001), with El Niño years that do not result in drought being associated with a positive IOD event (Ummenhofer et al. 2011). However, ENSO-IOD relationships have also weakened recently (Ham et al. 2017).

Ecological effects of ENSO include increased drought-related plant and animal mortality (Villalba & Vebelen 1998, Slik 2004) and fluctuations in ecosystem productivity and plant growth (Asner et al. 2000, Wigneron et al. 2020, Reis et al. 2024). The impact of ENSO on tree phenology has been studied extensively. One of the best examples of ENSO effects on reproductive phenology is the association of mass or “general” flowering and mast fruiting with droughts caused by El Niño events (Numata et al. 2003, Sakai et al. 2006, Brearley et al. 2007). However, La Niña episodes are also known to precipitate mass flowering through night-time temperature drops (Yasuda et al. 1999). The El Niño is also associated with an increase in flowering and fruiting at a number of sites where annual patterns of plant reproduction predominate (Wright et al. 1999, Chapman et al. 2018, Detto et al. 2018). Plant reproduction is expected to increase during El Niño episodes due to a rise in solar irradiance resulting from less cloud cover (Wright et al. 1999, Wright & Calderón 2006), which also allows plants to cope with the simultaneously occurring drought-like conditions (Detto et al. 2018). However, the validity of this pathway has not been examined through mechanistic modelling, such as path analysis. However, severe droughts caused by extreme El Niño could result in decreased plant reproduction owing to water stress (Wright & Calderón 2006). More generally, extreme warming adversely affects plant physiology by reducing the carbon balance (Dusenge et al. 2018).

In a prior study, we found that patterns of vegetative and reproductive phenology at a tropical forest site in the Indian Eastern Himalaya exhibited strong seasonality, with most species flowering once a year. Flowering and fruiting was highest during the warm dry season from March to May, with seasonal fruiting patterns based on seed dispersal mode. Daylength and solar irradiance were important as proximate cues for seasonal flowering and fruiting (Datta et al. 2025). However, even predictably seasonal phenological patterns can coexist with substantial inter-annual variation in leafing, flowering and fruiting (e.g. Chapman et al. 2018). A study of both inter-annual and seasonal patterns is required for a comprehensive understanding of tree phenology (e.g. Polansky & Robbins 2013), especially for more seasonal tropical forests, where the two kinds of patterns can be clearly separated.

Long-term trends in phenological patterns need to be studied concurrently with those of climate. The climate of the Northeast Indian region, which includes the Eastern Himalaya, is shaped heavily by the south-west monsoon from June to September, and exhibits strong intra-regional variation (Ravindranath et al. 2011, Dikshit & Dikshit 2014). Studies have been equivocal with respect to long-term monsoon rainfall trends in the region (Dash et al. 2012, Jain et al. 2013, Choudhury et al. 2019), even as mean and maximum temperatures have broadly increased (Dash et al. 2012, Jain et al. 2013). Monsoon rainfall trends in northeast India often exhibit distinct patterns from the rest of tropical South Asia, and severe El Niño events have either had no significant or even a positive effect on monsoon rainfall in the region (Kumar & Singh 2021).Therefore, ENSO-phenology relationships in the Eastern Himalaya, if mediated by climate variables, could show distinct characteristics compared to other tropical regions.

In this study, we examined long-term patterns in the intensity of tree reproductive phenology over a 14-year period in a tropical forest site in the Indian Eastern Himalaya. Inter-annual variation in flowering and fruiting, as distinct from seasonal variation, were examined at the community-level (i.e. for all 54 species that were monitored), for trees belonging to the principal seed dispersal modes (i.e. bird, mammal and mechanically-dispersed species). At the species level, we examined long-term trends for 36 species for which at least 10 trees were monitored. In addition, long-term trends in temperature, precipitation and solar irradiance were quantified. We expected to see an increasing trend in mean minimum and maximum temperatures across all seasons, in accordance with global (Forster et al. 2025) and regional (Dash et al. 2012, Jain et al. 2013) patterns, owing to anthropogenic climate change.

We sought to determine the climatic drivers of inter-annual variation in the intensity of flowering and fruiting for a 9-year subset (2011-2020) of the data, during which phenological and climate monitoring was relatively continuous. In particular, we examined whether ENSO was having a significant effect on inter-annual variation in reproductive intensity through the mediating effects of climate variables. Our approach incorporated lagged responses to ENSO, as these have been hypothesized (Wright et al. 1999) or empirically demonstrated (Chapman et al. 2018) to occur. Our first set of hypotheses pertained to the climatic pathways associated with inter-annual variation in reproductive phenology and these were: i) Solar irradiance was hypothesized to be the primary variable mediating ENSO-phenology relationships based on our previous study (Datta et al. 2025). As tree phenology in the site is strongly seasonal, we expected solar irradiance, the principal cue for reproductive phenology (and in particular, fruiting), to mediate inter-annual variation in flowering and fruiting, as hypothesized in Wright & Calderón (2006). ii) We had two competing hypotheses regarding ENSO effects: a) During El Niño events, if rainfall decreased, and thus, solar irradiance increased, then flowering and fruiting would also increase or b) If El Niño episodes resulted in a reduction in solar irradiance due to increased rainfall, then there would be reduced flowering and fruiting.

Our remaining hypotheses dealt with phenological and morphological or reproductive traits mediating species-specific variation in inter-annual phenological patterns. The traits pertained to a species’ intrinsic morphological or reproductive system such as the peak flowering and fruiting season, mode of reproduction, fruit/flower size, fruiting type and seed dispersal mode. We hypothesized that phenological responses to variations in solar irradiance would be more pronounced for species with larger flowers and fruits, as these are more energetically expensive to produce (based on models whereby per seed energetic investment is held constant, e.g. Stapanian 1982). We also expected responses to inter-annual variability in solar irradiance to be higher for mechanically-dispersed species, as these require dry conditions for fruit dehiscence and seed dispersal (van Schaik et al. 1993, Griz & Machado 2001).

## MATERIALS & METHODS

### Study Area

Pakke Tiger Reserve (862 km^2^; 92°36′–93°09′E and 26°54′–27°16′N) in western Arunachal Pradesh, India is located in the Eastern Himalaya Biodiversity Hotspot (Figure 1). Pakke has a tropical climate and is dominated by semi-evergreen forests, with major plant families being Lauraceae, Euphorbiaceae, Myrtaceae and Meliaceae in the lower elevation forests (Champion and Seth 1968, Datta & Rawat 2008, Page et al. 2022). The long-term average annual rainfall is 2500 mm. Pakke has a distinctly seasonal climate, with rainfall being concentrated during the southwest monsoon from June to September. The period from November to February consist of the cool dry, winter months, while that from March to May had warm and largely dry weather.

**Figure 1.**
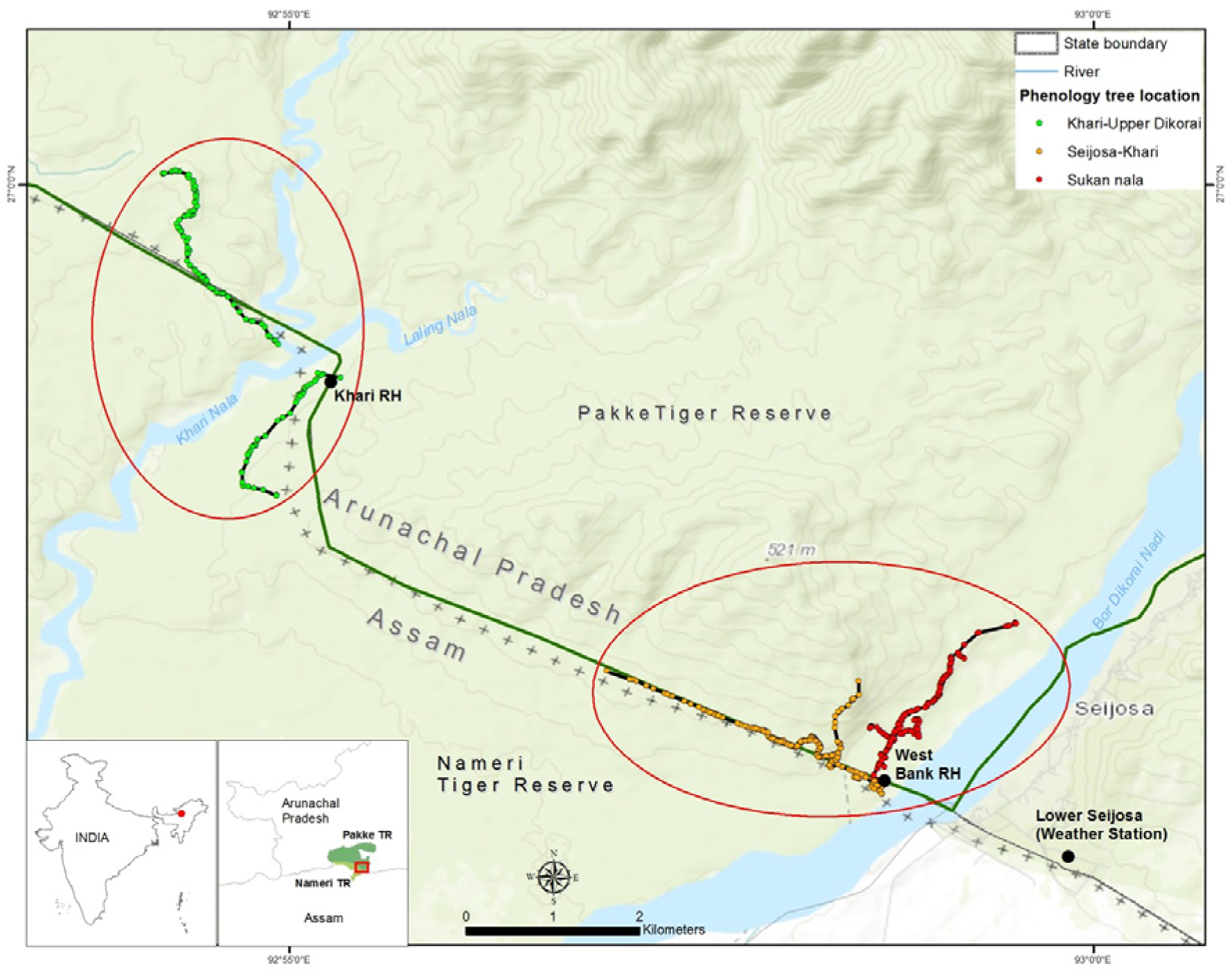
Map of trails used for phenology monitoring in Pakke Tiger Reserve, with the location in the state of Arunachal Pradesh in India highlighted.

The mean (± SD) maximum temperature in Pakke Tiger Reserve was 29.3°C ± 4.2°C and the mean minimum temperature was 18.3°C ± 4.7°C based on records from 1996-2000 ((Datta 2001). During the period from 2011 to 2023, the mean maximum temperature was 30.5°C and the mean minimum temperature was 18.6°C (Datta et al. 2025).

### Weather monitoring

A weather station (H21-USB data logger) was set up in April 2011 in Seijosa, at the edge of Pakke (1 km straight-line distance from the reserve) where temperature, rainfall, and solar irradiance are recorded every hour. We had complete weather data for all variables for 110 months between April 2011 and December 2024, with the remainder being lost to equipment malfunctions.

### Tree phenology monitoring

We monitored 716 reproductive trees of 54 tree species along four trails from January 2011 to December 2024 (see Datta et al. 2025 for more details). The number of trees monitored per species varied from one (four species) to 28 (for *Dillenia indica*). Two to four trained observers using binoculars assessed the presence of each phenophase in the tree canopies of tagged individuals. We recorded the presence/absence of flower buds, flowers, unripe fruit, ripe fruit, and scored as 1 if any or all of these reproductive phases are present and as 0 if absent. This provides a total count of the number of trees of each species in various phenophases in every month. The relative abundance of fruits on a tree was classified on an ordinal scale of 0-4, depending on the proportion of canopy covered by fruit (i.e. 0: no fruiting, 1: 0-25%, 2: 25-50%, 3: 50-75%, 4: 75-100%). For this paper, we are examining the long-term patterns in the reproductive phenology of 36 tree species for which at least 10 trees were monitored based on the monthly monitoring (25^th^–28^th^ of every month) from January 2011 to December 2024. These species include 7 mechanically dispersed, 8 mammal dispersed, 15 bird dispersed, and 6 species dispersed by both birds and mammals based on Datta (2001) and Datta and Rawat (2008).

## Analysis

### Characterizing trends in reproductive phenology and climate

We characterized long-term trends in flowering and fruiting over the study duration at the community level (i.e. across all 54 species) and for a subset of 36 tree species for which at least 10 trees were monitored. In addition, we characterized trends by pooling trees belonging to species for each of the principal seed dispersal modes (viz. bird-, mammal- and mechanically-dispersed).

Time series of the monthly proportion of trees in flower and the sums of fruiting scores were decomposed using the STL (Seasonal and Trend decomposition using LOESS) algorithm (Cleveland et al. 1990) over two periods, viz. 2011-2020 and 2021-2024. We treated these two periods separately owing to the 14-month long gap in phenological data collection in 2020-21. The robust STL decomposition algorithm has been used in a number of studies to examine phenological trends independent of seasonal variation or stochastic variation (e.g. Bhandari et al. 2012, Vogado et al. 2022). Gaps in monitoring during these periods were imputed through cyclical interpolation using the *na_seadec()* function of the *imputeTS* package (Moritz & Bartz-Bielstein 2017).

We decomposed the trend components of the time series that were Z-transformed for ease of interpretation of trends. Generalized Additive Modelling was used to fit cubic smoothed splines to these time series, with a basis dimension of 20 being used to model phenological trends in the former period (2011-2020) and 10 in the 2021-2024 period. We feel that this represents an adequate trade-off between model accuracy and overfitting. In order to remove the high degree of temporal autocorrelation characteristic of phenological time series, additive autoregressive terms were progressively incorporated till the Ljung-Box test of temporal autocorrelation returned a non-significant result for the residuals (Ljung & Box 1978). We used tests of significance of the F statistic and the effective degrees of freedom of the additive model to infer the existence of nonlinear trends, which were visually examined using plots of the smooth functions. However, trends for the period 2021-2024 should be interpreted cautiously, as the time series during this period was short. Generalized Additive Modelling has been used to study long-term phenological trends in Uganda (Polansky & Robbins 2013) and Australia (Vogado et al. 2022).

Trends in climate variables were also assessed using cubic smoothing splines (as in Vogado et al. 2022), and this was done only for the period from 2011-2020, as climate data was sparse in 2021-2024. Cyclic interpolation was used to fill gaps in climate data in the period 2011-2020, as with phenology data. In addition, we used linear regression to examine for significant trends in the seasonal means of temperature, precipitation and solar irradiance. The following seasons were delineated: i) March to May: warm dry (summer), ii) June to September: wet season (southwest monsoon) iii) November to February: cool dry (winter), and October was considered to be a transitional month between the southwest monsoon and the cool dry months. Linear modelling was carried out on climate data collected between April 2011 and July 2022. The R package ‘*mgcv’* was used to carry out Generalized Additive Modelling (Wood 2011).

### Examining climate-phenology relationships

We used Generalized Linear Mixed Models (GLMMs) using the *glmmTMB* package in R (Bolker et al. 2009, Brooks et al. 2017) to examine the effects of ENSO, temperature, precipitation and solar irradiance on inter-annual variation in the intensity of tree reproductive phenology. We utilized time series of species-level phenology and climate variables between January 2011 and February 2020 for each of the 36 tree species. We preferred GLMMs to GAMs for modelling climate-phenology relationships, as GLMMs allow for easy interpretation of parameter effects.

For flowering, the response variable consisted of species-specific time series of the number of trees in flower in relation to the number that did not, and, was therefore modelled using beta-binomial distribution. On the other hand, the fruiting response variable was the sum of intensity scores (on a scale from 0-4), which was assumed to follow a negative binomial distribution, as it was an integer variable. In each GLMM, the month of the year was included as a covariate, in order to ensure that the residuals did not incorporate seasonal variation. The proportion of trees in flower and the fruiting score in the preceding month was used to model an autoregressive structure (i.e. AR(1)) in flowering and fruiting models respectively. For flowering models, there was a significant correlation between residuals lagged by a year, and therefore, the 12-month lagged variable of the proportion of trees in flower was included as a covariate as well.

For both flowering and fruiting, we created two sets of models: one incorporating ENSO effects and the other representing inter-annual variation in climate variables related to temperature, precipitation and solar irradiance. The effects of ENSO were quantified by the Multivariate ENSO Index (MEI), a robust and comprehensive basin-wide measure (Wolter & Timlin 1998) that is widely used in phenological research (e.g. Brown et al. 2010, Robichaud & Comtois 2017, Chapman et al. 2018). In the climate model set, predictors consisted of the de-seasonalized residuals of five climate variables: i) mean maximum temperature ii) minimum temperature iii) monthly mean of the daily maximum solar irradiance iv) total monthly rainfall and v) the proportion of rainy days in a month. Seasonal patterns in these climatic variables were removed using a linear model in which month was used as a factor variable, and the residuals were used as predictors. This de-seasonalization process is conceptually similar to that in Polansky & Boesch (2013), but we used month as a factor variable instead of a spline, to account for complex and noisy seasonal patterns which splines may fail to accommodate.

As plant phenology is known to exhibit lagged responses to climatic perturbations (e.g. Sherry et al. 2011, He et al. 2024), we looked at lags in flowering and fruiting responses across 0-12 months corresponding to MEI and for each climate variable. This was done using a two-step process. First, we used cross-correlation functions (CCF) to determine the extent to which flowering and fruiting lagged behind MEI and each climate variable. For this purpose, phenological and climatic time series were decomposed using STL, and the seasonal component was removed. MEI was not decomposed, as it reflects temperature anomalies and is thus independent of seasonal patterns. Thereafter, for each MEI-phenology and climate-phenology relationships, only those monthly lags in phenological responses were taken into consideration for which the absolute value of the cross correlation function was > 0.2, as these were felt to indicate potentially significant relationships. The CCF value threshold was deliberately kept low, as the objective of running CCFs was to screen for potential lags and reduce the number of lagged responses being modelled by GLMMs, while the GLMMs themselves would be used to assess statistical significance of effect sizes.

Thereafter, we created two sets of GLMMs for either phenophase, one corresponding to the effects of MEI and the other related to climate variables. In each set, individual GLMMs were created corresponding to each of the lagged responses screened using the process outlined in the previous section. The climatic predictors corresponding to temperature, precipitation and solar irradiance were the residuals obtained from a linear model with month as a factor variable, in order to ensure that we were only modelling phenological responses to inter-annual climatic variation.

Each GLMM accommodated only a single lagged response in order to avoid multicollinearity and to pinpoint the specific extent of the lag for which climate-phenology relationships were most significant. Inferences based on effect strength or cross correlations across monthly lags have been used in modelling long-term ENSO-phenology and ENSO-climate relationships (e.g. Roy & Sparks 2000, Chapman et al. 2018). The 95% confidence intervals of climate predictors were used to infer significance of effect sizes, and the Akaike Information Criterion optimized for small sample sizes (Burnham & Anderson 2002) was used for model comparisons across GLMMs in each model set. If predictor effects were significant for the best-supported models in both sets, Structural Equation Modelling was used to infer whether ENSO effects were being mediated by climate variables, i.e. to determine if there was adequate support for an MEI→climate→ phenology pathway (Shipley 2000). The constituent models of the pathway involved the most parsimonious MEI and climate models. All variables were z-transformed to facilitate comparisons of effect sizes and model convergence, and species identity was used as a random effect.

In order to determine whether species-specific responses were being mediated through traits, we used interactive models of traits (morphological, reproductive and phenological) and climate/ENSO variables represented by the respective best-supported models. Traits are represented in the following table (Table 1). Diagnostic tests of GLMMs were carried out using the *‘DHARMa’* package in R (Hartig 2025). The modelling pathway is represented as an infographic in Figure 2.

**Figure 2.**
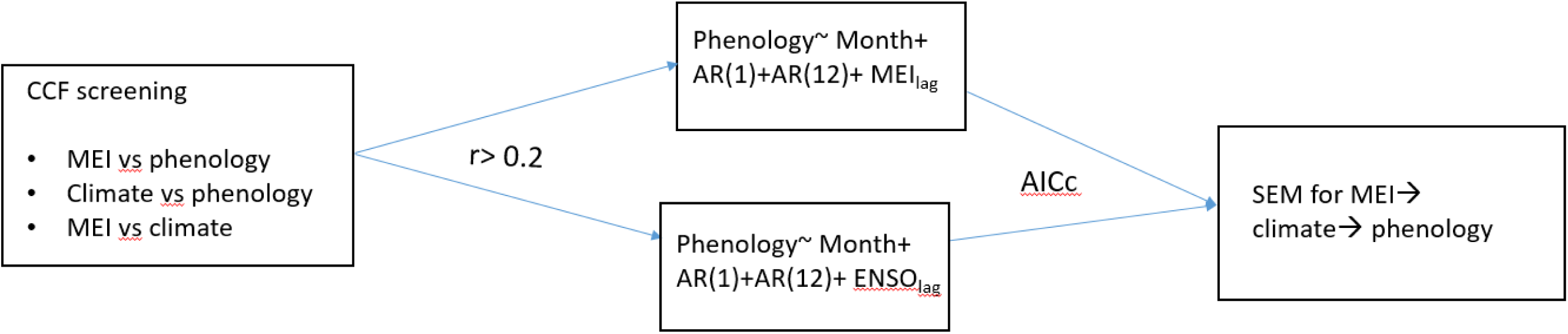
Infographic highlighting the modelling pathway used for modelling effects of Multivariate ENSO index (MEI) and climate on inter-annual patterns in tree flowering and fruiting in Pakke Tiger Reserve, Arunachal Pradesh, India. CCF: cross correlation function, r: value of CCF, ENSO: El Niño Southern Oscillation, AICc: Akaike Information Criterion corrected for small sample size, SEM: Structural equation modelling.

**Table 1.**
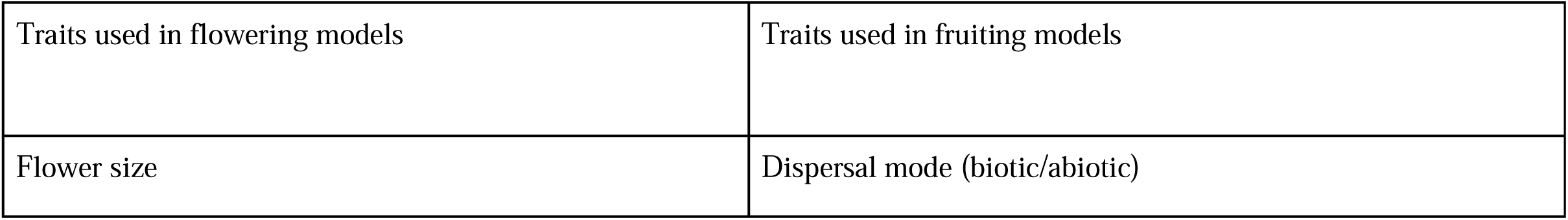
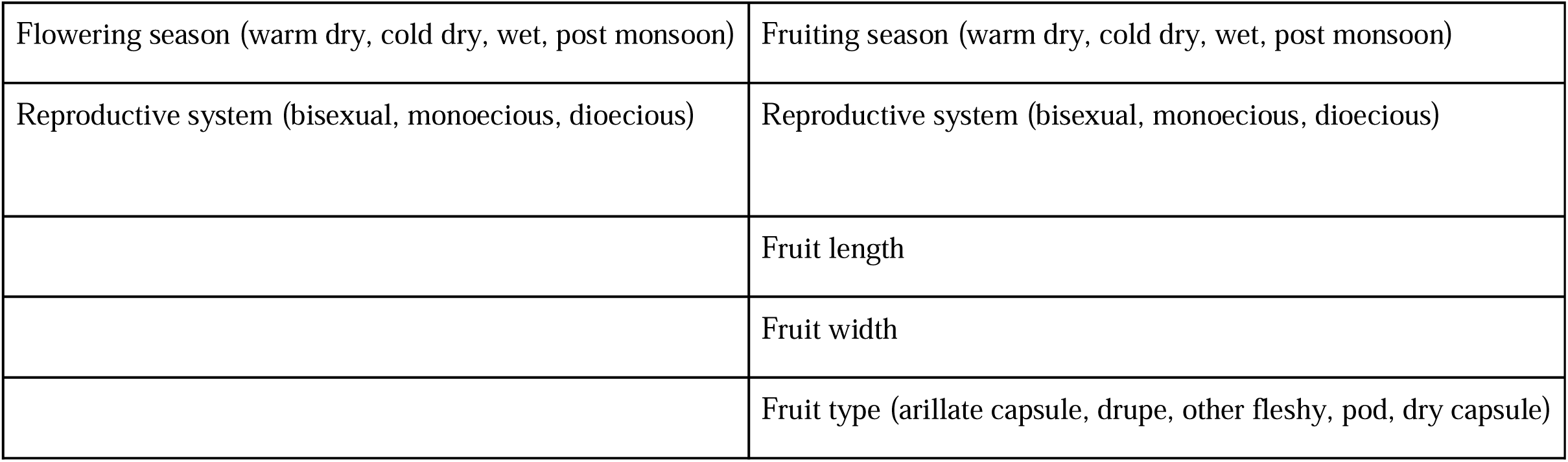
List of all morphological, reproductive and phenological traits used in modelling species-specific responses to flowering and fruiting in the study area.

|  |  |
| --- | --- |
| Traits used in flowering models | Traits used in fruiting models |
| Flower size | Dispersal mode (biotic/abiotic) |

|  |  |
| --- | --- |
| Flowering season (warm dry, cold dry, wet, post monsoon) | Fruiting season (warm dry, cold dry, wet, post monsoon) |
| Reproductive system (bisexual, monoecious, dioecious) | Reproductive system (bisexual, monoecious, dioecious) |
|  | Fruit length |
|  | Fruit width |
|  | Fruit type (arillate capsule, drupe, other fleshy, pod, dry capsule) |

## RESULTS

### Trend in long-term flowering during the study period

In the period 2011-2020, a significant nonlinear trend in flowering was observed at the community-level (i.e. considering all monitored trees as a whole, edf = 3.424, F = 3.986, p = 0.005) and for each of the principal seed dispersal modes (bird-dispersed: edf = 3.19, F = 3.032, p = 0.022, mammal-dispersed: edf = 2.563, F = 3.451, p = 0.017, mechanically-dispersed: edf=9.185, F = 3.955, p <0.0001). We also observed significant nonlinear flowering trends for 17 out of 35 species (i.e. excluding *Ficus drupacea*, a fig which has no visible flowers) in 2011-2020 (Table S1), while two others (*Endospermum chinense* and *Horsfieldia kingii*) showed significant trends that were effectively linear (Table S1). Community-level flowering showed an approximately monotonic increase in the period 2011-2016, with a slight reduction thereafter (Figure 3). Flowering patterns exhibiting a similar peak were also shown by bird- and mammal-dispersed trees, while for mechanically-dispersed trees (Figure 4), there was a much stronger flowering increase in the period 2015-2016. For 14 species (viz. *Actinodaphne obovata, Beilschmiedia* sp., *Choerospondias axillaris, Chukrasia tabularis, Cryptocarya* sp.*, Dysoxylum cauliflorum, Dysoxylum procerum, Gynocardia odorata, Knema angustifolia, Magnolia hodgsonii, Pterygota alata, Picrasma javanica, Stereospermum tetragonum, Sterculia villosa*) (Fig. S1), flowering peaked around 2015-16, with a small decline or mostly stable flowering intensity in 2016-2019. For 7 species (viz. *Baccaurea ramiflora, Beilschmiedia* sp., *C. tabularis*, *E. chinense*, *H. kingii*, *S. tetragonum*, *P. alata*), flowering appeared to be higher at the end of the first monitoring period than at the beginning. Flowering was lower in 2019-20 than in 2011 only for *Livistona jenkinsiana* was flowering lower in 2019-20 than in 2011.

**Figure 3.**
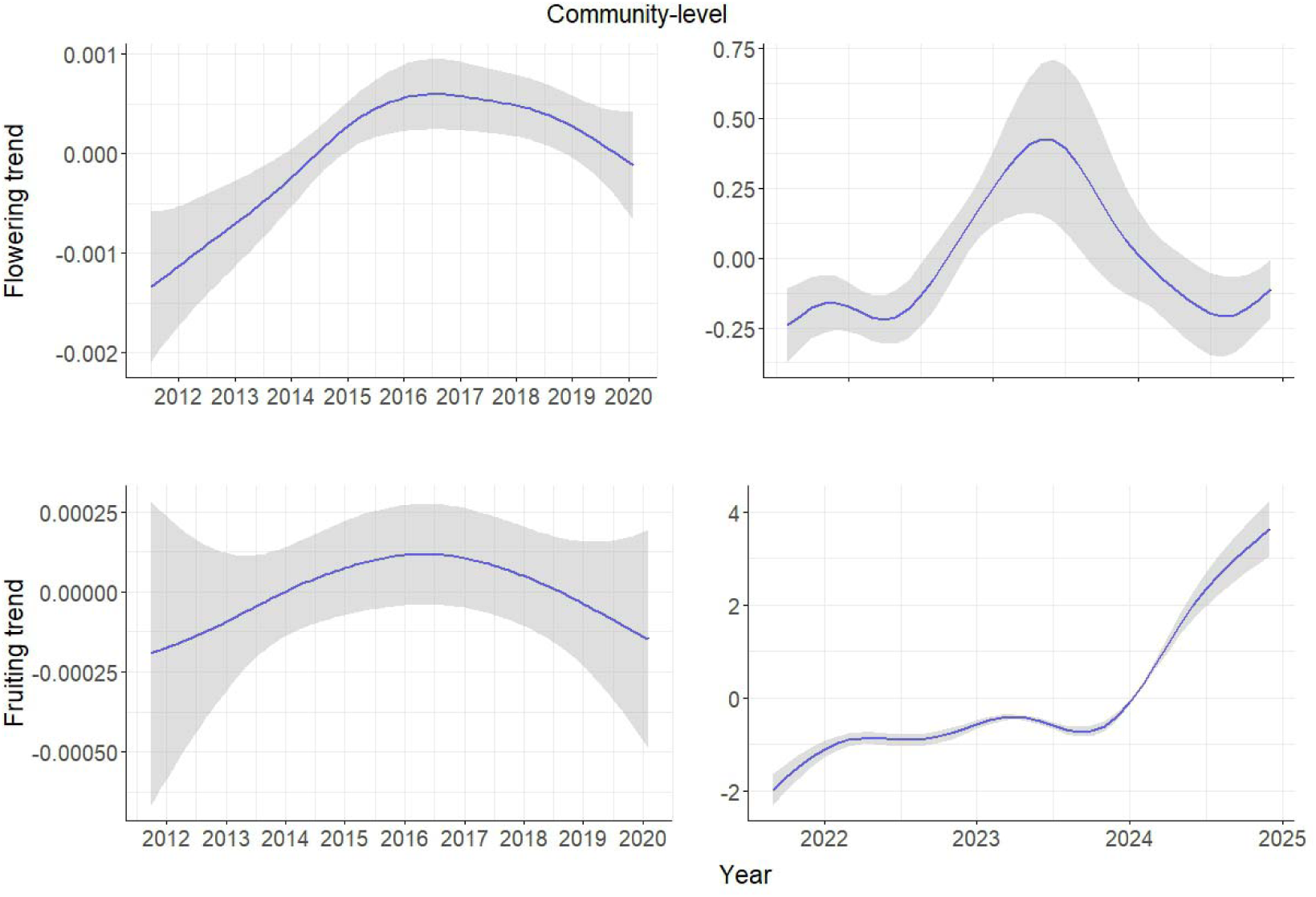
Generalized Additive Modelling (GAM) smooth functions of the trend in community-level phenology in the 2011-2024 period.

**Figure 4.**
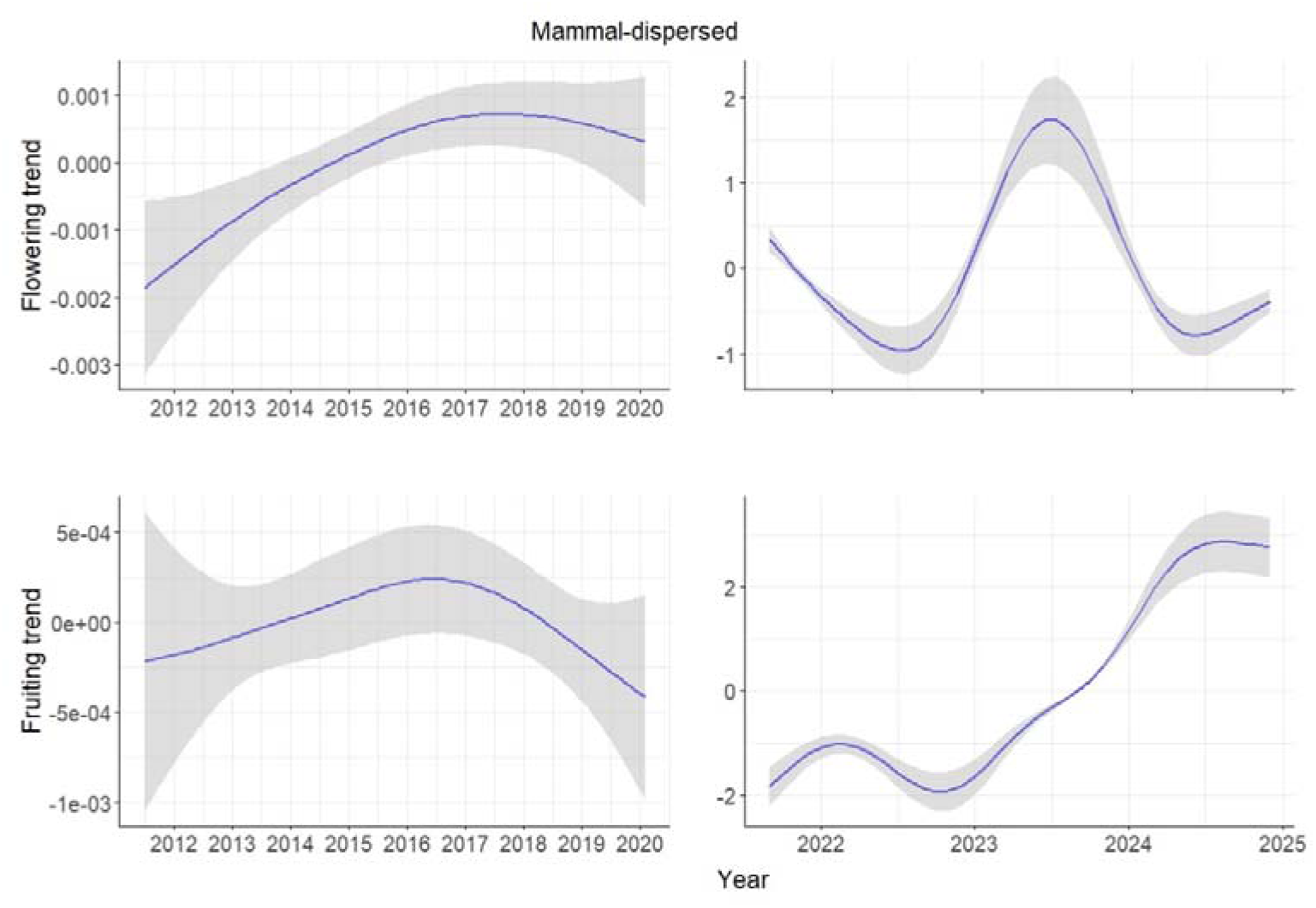
Generalized Additive Model (GAM) smooth function indicating the trend in flowering and fruiting of trees belonging to mammal-dispersed species between 2011 and 2024 in the Pakke Tiger Reserve, Arunachal Pradesh, India.

**Figure 5.**
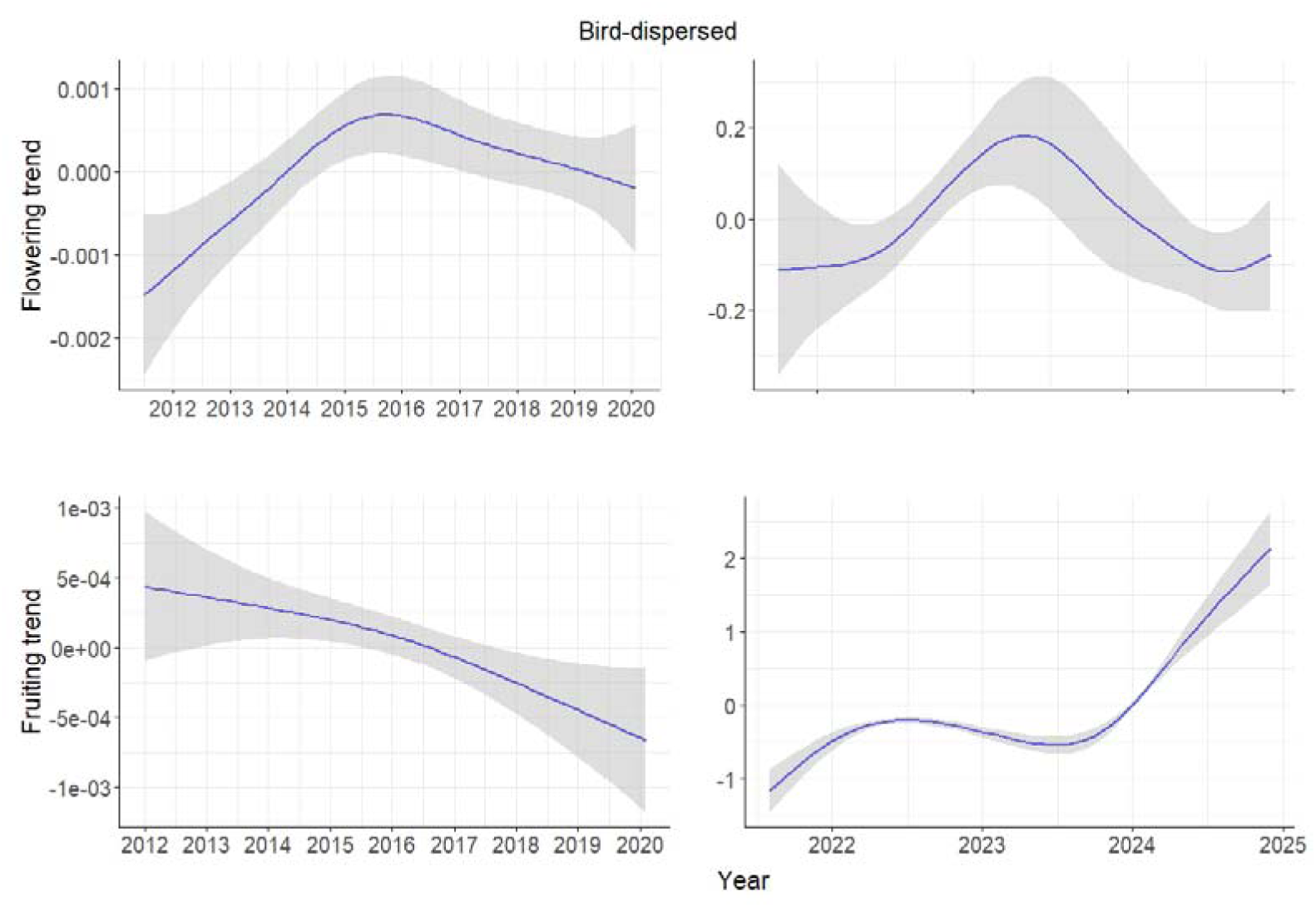
Generalized Additive Model (GAM) smooth function indicating the trend in flowering and fruiting of trees belonging to bird-dispersed species between 2011 and 2024 in the Pakke Tiger Reserve, Arunachal Pradesh, India.

During the latter monitoring period (i.e. from April 2021-December 2024), flowering trends were significantly nonlinear at the community-level and for 27 out of 35 species (Table S1). At the community-level (Figure 3) and for 12 of the species, flowering trends during this period were distinctly unimodal, peaking around mid-2023 (Fig. S1). Qualitative examination of the maximum percentage of trees in flower in any year corroborated this trend, for e.g. *C. axillaris* had all monitored trees flowering in 2016 and 2023, but this was only 7-13% between 2011 and 2013. Across all species, the maximum percentage of species in flower at any month was highest in 2023 (25%) followed by 2016 (22%), and the years with lowest flowering were 2022 and 2024 (both 11%).

### Trends in long-term fruiting during the study period

In the 2011-2020 period, the trend in long-term fruiting patterns at the community level (Figure 1) or for mammal-dispersed trees (Figure 4) was not statistically significant (Table S1). However, bird-dispersed trees showed a significant, broadly declining trend over this period (edf = 1.689, F = 3.634, p = 0.03), whereas mechanically-dispersed trees showed a significant bimodal trend, with peaks in 2014 and 2018 (edf = 14.670, F = 7.503, p < 0.0001) (Fig 4 & 5).

Significant nonlinear trends in fruiting in the period 2011-2020 were exhibited by 21 species, while two others (viz. *Chisocheton cumingianus* and *Zanthoxylum rhetsa*) showed roughly linear trends (Figure S1 & Table S1)

Visual inspections of the spline functions indicated that for 8 species (viz. *Alstonia scholaris*, *B. ramiflora*, *H. kingii, Monoon simiarum, M. hodgsonii, K. angustifolia, Prunus ceylanica* and *Z. rhetsa*), the fruiting in 2019-2020 was lower than that at the start of the sampling period (viz. 2011), with substantial periods of declining fruiting (Fig 7, 8 & S1). This appeared to be corroborated by qualitative examinations of fruiting data. For instance, *B. ramiflora* showed no fruiting between 2015 and 2019, whereas, in *M. hodgsonii*, the maximum percentage of trees in fruit at any month of the year declined from 90% in 2011 to just 25% in 2019, and continued to decline even further thereafter, being just 11.76% in 2024. Fruiting was higher at the end of the first monitoring period, compared to 2011 only for *S. tetragonum*.

Fruiting trends in the period from May 2021 to December 2024 should be interpreted with caution owing to the relatively high effective degrees of freedom of GAM smoothers (edf ≈ 7-9) indicating potential overfitting. A significant nonlinear fruiting trend was observed at the community level and for all dispersal modes during this period, with an especially distinct increase over the year 2024 (Figures 3-6). In addition, 34 out of 36 species showed significantly nonlinear trends in fruiting. Visual examination of spline functions indicated fruiting increased over this period for 23 of these species (Fig. S1). These included three species (viz. *K. angustifolia*, *Z. rhetsa* and *P. ceylanica*) that had declining trends in the first period.

**Figure 6.**
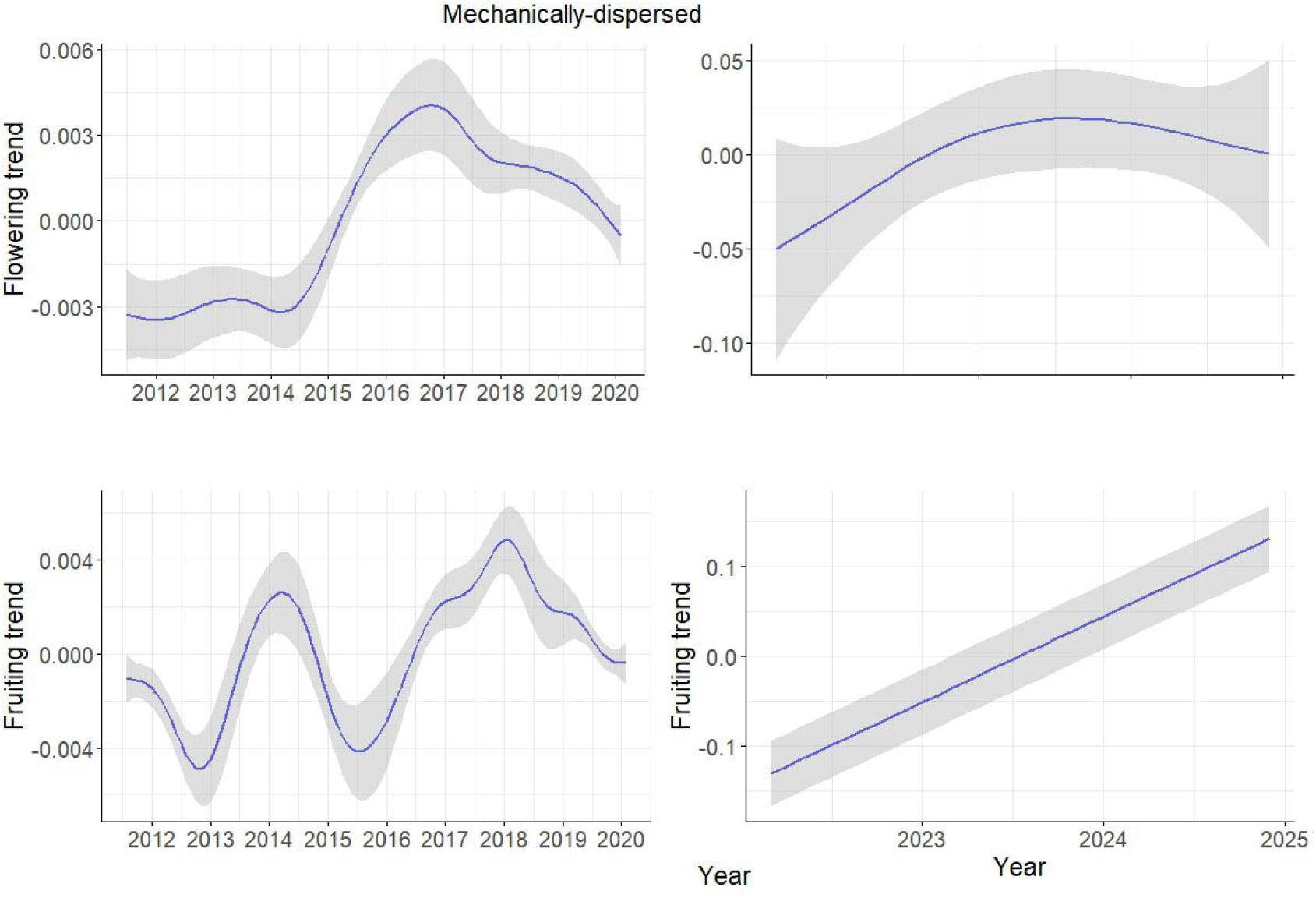
Generalized Additive Model (GAM) smooth function indicating the trend in flowering and fruiting of trees belonging to mechanically-dispersed species between 2011 and 2024 in the Pakke Tiger Reserve, Arunachal Pradesh, India.

**Figure 7.**
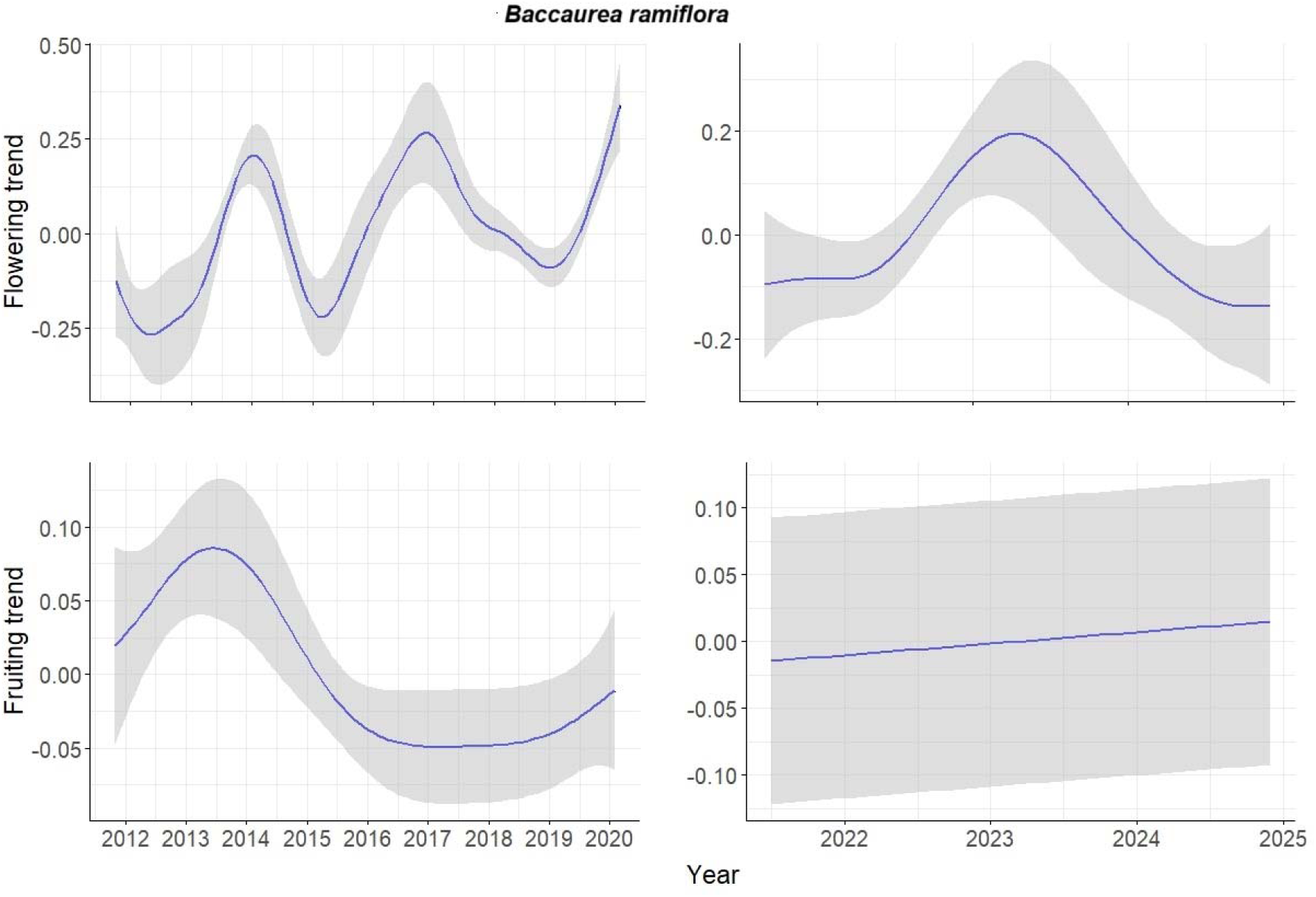
Generalized Additive Model (GAM) smooth function indicating the trend in flowering and fruiting of *Baccaurea ramiflora* between 2011 and 2024 in the study area.

**Figure 8.**
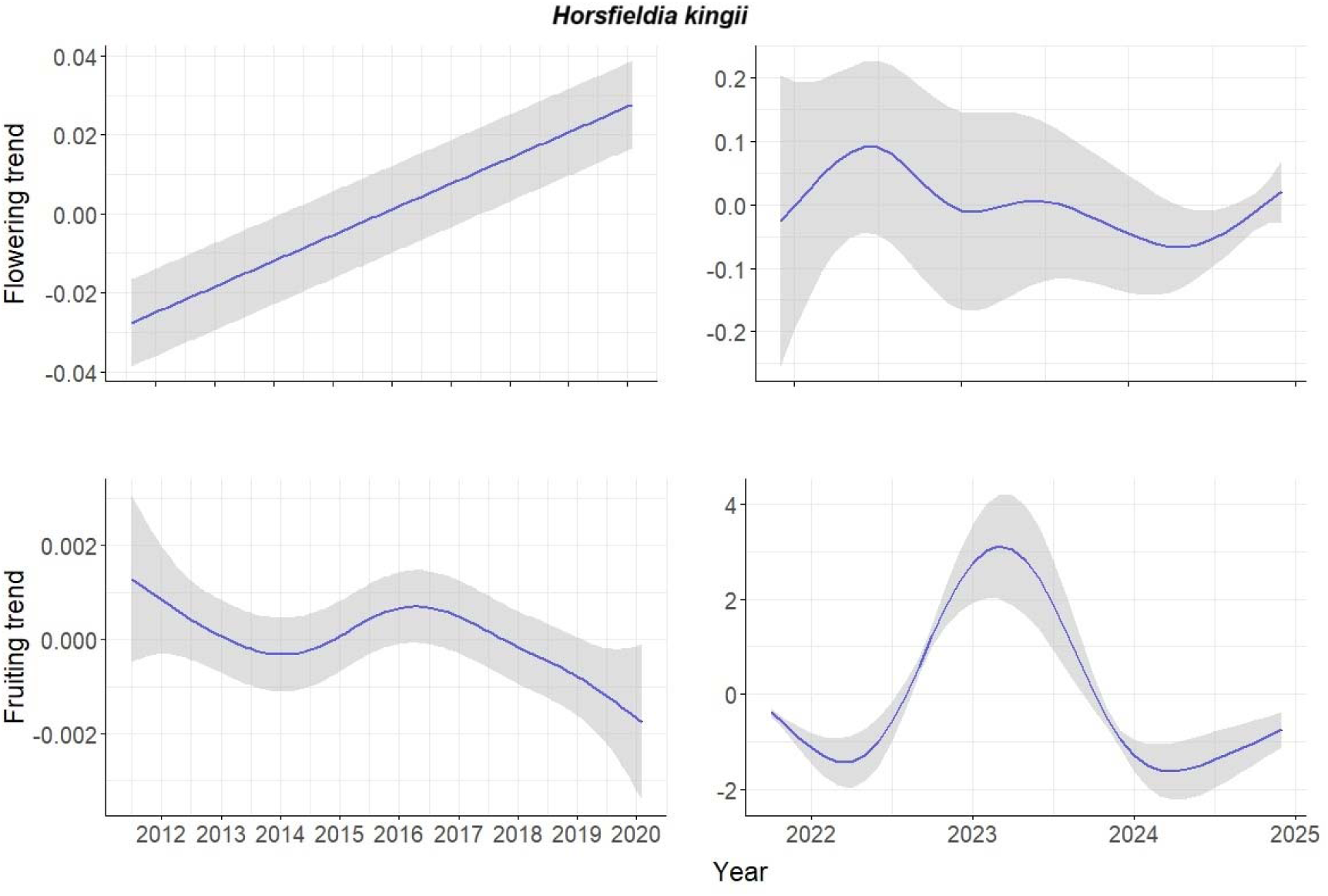
Generalized Additive Model (GAM) smooth function indicating the trend in flowering and fruiting of *Horsfieldia kingii* between 2011 and 2024 in the study area.

### Trends in climate variables

Generalized Additive Modelling indicated substantial nonlinear trends for mean minimum temperature (edf = 15.33, F = 7.96, p < 0.0001) and total rainfall (edf =13.65, F = 5.707, p <0.0001) and the monthly proportion of rainy days (edf =16.31, F = 10.26, p <0.0001) in the period 2011-2020. Plots of the corresponding smooth variables indicate a general increase in minimum temperature and precipitation over this period. (Figure 9). Linear models indicated a decline in mean solar irradiance during the warm dry season ((β = -21.718 (-35.725, -5.711), p = 0.015, R^2^ = 0.54)). The wet season showed an increase in both mean maximum (β = 0.191 (0.040, 0.343), p = 0.012, R^2^ = 0.42) and mean minimum temperatures ((β = 0.080 (0.051, 0.109), p <0.001, R^2^ = 0.79)). None of the seasons showed a significant trend in either mean monthly precipitation or the monthly proportion of rainy days (Table S2).

**Figure 9.**
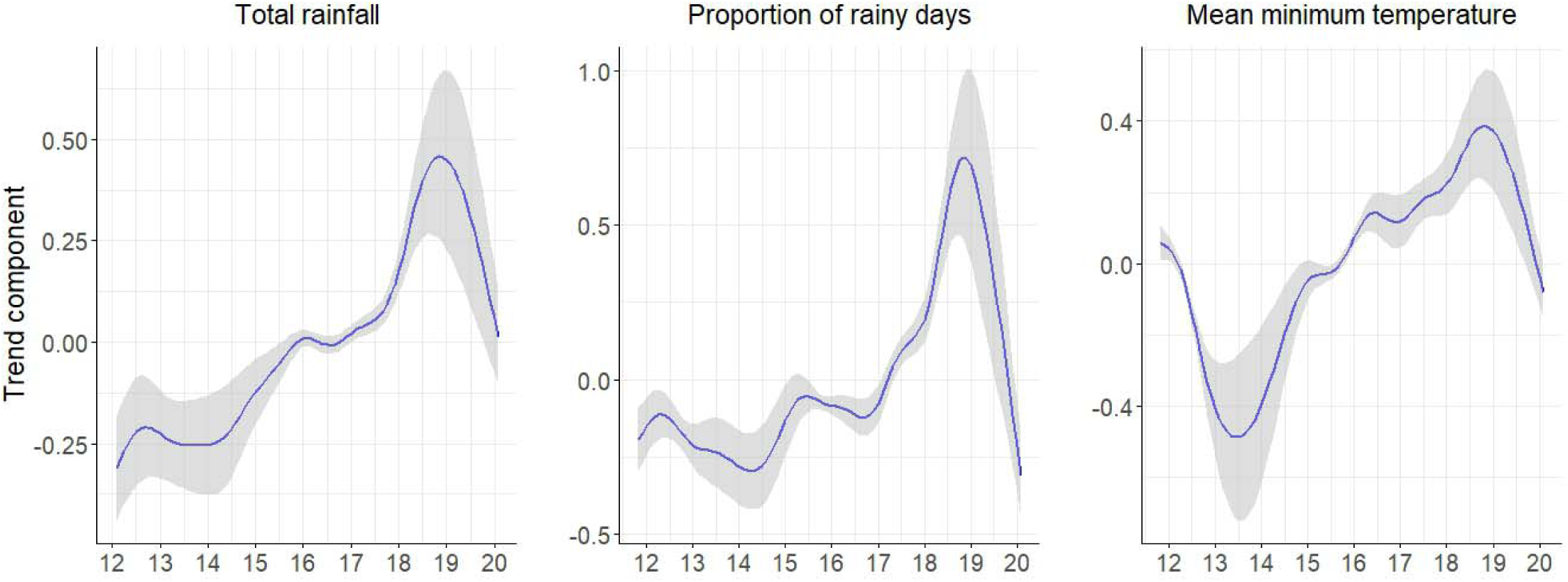
Generalized Additive Models (GAM) smoothing functions for total rainfall, monthly proportion of rainy days and mean minimum temperature in the period 2011-2020.

The trend in the MEI in 2011-2024 indicated that 2011-2012 March, 2017 July-2018 June, 2020 June-2023 March and July-December 2024 were periods of negative ENSO anomalies corresponding to La Niña, while the periods from May 2015- May 2016 and July 2023-March 2024 were associated with El Niño (Figure S2).

### EFFECTS OF ENSO, CLIMATE AND TRAITS ON LONG-TERM FLOWERING AND FRUITING

Cross-correlation function plots appeared to indicate a strong positive association between MEI and flowering, as well as between MEI and monthly rainfall for lags of up to two months. The correlation between MEI and solar irradiance over this period appeared to be negative (Figure S3). Based on values of the cross correlation coefficient > ∼0.2, the following predictors were chosen for the flowering and fruiting models (Table 2):

**Table 2.** List of MEI or climate variables to which lagged phenological responses were modelled in GLMMs. The lags appended to each variable name refer to the corresponding number of months by which flowering or fruiting was considered to be lagged in relation to these variables.

| Phenophase | Lag phenological responses for MEI variables | Lagged phenological responses for climate variables |
| --- | --- | --- |
| Flowering | MEI lag0 - lag7 | Solar irradiance: lags 0, 2, 3<br>Total rain: lag 4<br>Monthly rainy days: lag 0 |

|  |  |  |
| --- | --- | --- |
| | | (For temperature variables, CCF values for all lags $\ll 0.2$) |
| Fruiting | MEI lag 0, lag 1 | Mean maximum: lag 0<br>Solar irradiance: lags 0 and lag 8<br>Total rain: lag 8<br>Rainy days: lag 0 and lag 8<br>(For mean minimum, no lagged responses modelled as $r \ll 0.2$ for all cases) |

The best-supported ENSO model for flowering incorporated the effect of MEI without any monthly lags, which explained 40% of the variation in flowering (conditional Nakagawa & Schielzeth’s R^2^=0.40) (Table 3). MEI had a significantly positive association with flowering (β_MEIlag0_= 0.253 (0.170, 0.336)).The other ENSO models had only negligible AICc weight (Table S3). Among the climate models, the best-supported flowering model consisted of the effect of mean maximum temperature without any monthly lags (Table S4). However, the effect of this predictor was not significant (β_mean maximum temperature lag0_= -0.051 (-0.124, 0.023)).

**Table 3.** Output of the Generalized Linear Mixed Model (GLMM) incorporating the effect of Multivariate ENSO Index (MEI) without lags on long-term flowering in the study area. Estimates corresponding to the levels of the factor variable month are not provided for the purpose of brevity.

| Parameter | Beta coefficient (95% CI) | p value |
| --- | --- | --- |
| MEI (without lags) | 0.253 (0.170, 0.336) | <0.0001 |
| AR(1) | 1.899 (1.605, 2.194) | <0.0001 |
| AR(12) | 2.935 (02.644, 3.226) | <0.0001 |
| SD of random intercept | 0.344 (0.243, 0.486) |  |

No morphological trait was found to mediate species-specific flowering responses to ENSO (Table S5). However, the increase in flowering during El Niño was lower for species whose flowering peak was in the wet season, compared to those in the warm dry season (β_wet season*MEI_lag0_= - 0.193 (-0.381, -0.006). Species whose peak month of flowering was in the cool dry or post monsoon period did not show differing ENSO responses in relation to summer flowering species (Table 4).

**Table 4.** Output of the Generalized Linear Mixed Model (GLMM) incorporating the interactive effect of MEI and flowering season on long-term flowering in the study area. Estimates corresponding to the levels of the factor variable month are not provided for the purpose of brevity.

| Parameter | Beta coefficient(95% CI) | p value |
| --- | --- | --- |
| MEI (without lags) | 0.296 (0.180, 0.411) | <0.0001 |
| Wet | 0.502 (0.211, 0.794) | <0.0001 |

|  |  |  |
| --- | --- | --- |
| Post monsoon | 0.402 (-0.132, 0.936) | 0.140 |
| Cool dry | 0.214 (-0.194, 6.623) | 0.304 |
| MEI (without lags)* Wet | -0.193 (-0.381, -0.006) | 0.008 |
| MEI (without lags)* Post monsoon | -0.174 (-0.565, 0.217) | 0.384 |
| MEI (without lags)* Cool dry | 0.215 (-0.038, 0.468) | 0.096 |
| AR(1) | 1.908 (1.613, 2.202) | <0.0001 |
| AR(12) | 2.981 (2.688, 3.274) | <0.0001 |
| SD of random intercept | 0.277 (0.183, 0.410) |  |

For fruiting, the most parsimonious ENSO model represented the immediate, unlagged effect of MEI, while the only other model, which represented the lagged response by 1 month received non-negligible support, and this represented the lagged response by one month. Both models received an almost equal degree of support (AICc weight=0.497 vs 0.502). Neither variable had a statistically significant effect (β_MEI lag 0_= -0.009 (-0.132, 0.114) and β_MEI lag 1_= -0.002 (-0.129, 0.125)).The best supported climate model consisted of the 3-month lagged effect of solar irradiance (conditional Nakagawa & Schielzeth’s R^2^=0.32) (Table S6). Independent of the mediating effect of species-specific traits, inter-annual variation in solar irradiance and fruiting were positively associated (β_solar irradiance lag 3_=0.163 (0.044, 0.282)) (Table 4). Species with a fruiting peak in October showed a stronger positive response to inter-annual increases in solar irradiance (β_post monsoon*solar irradiance lag 3_=0.336 (0.022, 0.651), conditional Nakagawa & Schielzeth’s R^2^=0.33 for solar irradiance*fruiting season model), (Table 5). None of the morphological traits showed significant associations with inter-annual fluctuations in climatic variables (Table S7). Since at least one of the links in the MEI-climate-phenology casual pathway did not have statistically significant support for both flowering and fruiting, we did not perform Structural Equation Modelling.

**Table 5.** Output of the Generalized Linear Mixed Model (GLMM) incorporating the effect of solar irradiance on long-term fruiting in the study area. Monthly estimates of fruiting, used for removing seasonal patterns, are not provided for the purpose of brevity.

| Parameter | Beta coefficient(95% CI) | p value |
| --- | --- | --- |
| Solar irradiance (3 month lagged response) | 0.163 (0.044, 0.282) | 0.007 (<0.05) |
| Number of trees | 0.020 (-0.038, 0.078) | 0.503 |
| AR(1) | 0.175 (0.126, 0.225) | <0.0001 |
| SD of random intercept | 0.730 (0.524, 1.017) |  |

**Table 6.** Output of the Generalized Linear Mixed Model (GLMM) incorporating the interactive effect of MEI and fruiting season on long-term fruiting in the study area. Estimates corresponding to the levels of the factor variable month are not provided for the purpose of brevity.

| Parameter | Beta coefficient(95%CI) | p value |
| --- | --- | --- |
| Solar irradiance (3 month lagged response) | 0.095 (-0.104, 0.294) | 0.351 |

|  |  |  |
| --- | --- | --- |
| Number of trees | 0.025 (-0.031, 0.081) | 0.384 |
| AR(1) | 0.178 (0.129, 0.228) | <0.0001 |
| Cool dry | 0.781 (0.083, 1.479) | 0.029 (<0.05) |
| Warm dry | 0.014 (-0.591, 0.620) | 0.963 |
| Post monsoon | 0.713 (-0.056, 1.482) | 0.069 |
| Solar irradiance (3 month lagged response)* Cool dry | -0.054 (-0.366, 0.258) | 0.737 |
| Solar irradiance (3 month lagged response) * Warm dry | -0.014 (-0.358, 0.330) | 0.937 |
| Solar irradiance (3 month lagged response) * Post monsoon | 0.336 (0.022, 0.651) | 0.036 (<0.05) |
| SD of random intercept | 0.625 (0.437, 0.895) |  |

### ENSO effects on temperature, precipitation and solar irradiance

We found some support of an increase in total monthly precipitation (β_MEI lag 1_= 0.190 (-0.012, 0.392) and a reduction in solar irradiance (β_MEI lag 1_= -0.200 (-0.402, 0.002) about one month after an El Niño event (Table S8). For both these climate variables, the model representing the one month lagged response in relation to ENSO received maximum support. However, confidence intervals of the beta coefficient still overlapped zero, although only marginally, and the p values testing for a significant deviation of the beta coefficient from zero were 0.052 and 0.065 respectively (Table S8). ENSO did not appear to have an effect on temperature or the proportion of rainy days in a month. These findings seemed to be in accordance with the values of the cross-correlation function between MEI and climate variables (Figure S3).

## DISCUSSION

### Long-term phenological trends in flowering and fruiting

A substantial proportion of species showed significant nonlinear trends in reproductive phenology over the study period. Inter-annual variation in flowering and fruiting intensity is typically nonlinear in the seasonal tropics (Chapman et al. 2005, 2018, Mendoza et al. 2018, Polansky & Robbins 2013, Vogado et al. 2022), and our results adhered to this overall pattern.

Visual examinations of the spline functions appeared to indicate that periods of peak flowering coincided with El Niño episodes. At the community level and for 10 species that had significant nonlinear trends, flowering appeared to peak in 2015-2016 and again in 2023, which coincided with the two strongest episodes of El Niño during the period of the study. Likewise, qualitative examinations of the maximum percentage of trees in flower in any month indicated that flowering was lowest during the La Niña years of 2022 and 2024. Long-term fruiting patterns appeared to be unrelated to El Niño/La Niña episodes. Among the species that showed a declining fruiting trend in 2011-2020, all but one (viz. *A. scholaris*) were animal-dispersed. In addition, bird-dispersed species as a group showed a significant fruiting decline during this period. On the other hand, positive fruiting trends appeared to predominate for most species in 2021-2024. However, for some animal-dispersed species such as *Magnolia hodgsonii* and the locally abundant *Monoon simiarum*, the maximum percentage of trees in fruit remained much lower in 2024 than in 2011. Declines in fruit availability can adversely affect populations of large-bodied frugivores (Bush et al. 2020), necessitating the analysis of long-term fruiting trends based on seed dispersal mode. Thus, although a conclusive statement cannot be made about fruiting trends in the study area, it is vital to focus monitoring on select animal-dispersed species that are showing fruiting declines over substantial periods. Between 2011 and 2020, positive trends in flowering outnumbered negative ones while more species had negative fruiting trends. For two species with significant nonlinear trends (viz. *H. kingii* and *B. ramiflora*), flowering appeared to show an increase during this period while fruiting declined. Few studies have reported concurrently on trends of flowering and fruiting from the same site. Divergence between long-term flowering and fruiting patterns have been reported in Panama (Wright & Calderón 2006) and in Australian lianas (Vogado et al. 2022). The process of ripe fruit formation involves pollination, fertilization, fruit development and ripening, all of which are known to be susceptible to climatic extremes associated with climate change (Hoiss et al. 2015, Menzel 2019, Haokip et al. 2020).. In particular, ongoing declines in pollinator populations and phenological mismatch due to climate change (Kudo & Ida 2013, Soroye et al. 2020) and landuse conversion are well studied, and thus the role of pollination failure in the divergence between flowering and fruiting should be examined.

We emphasize here that reduced or absent flowering or fruiting in some years is unlikely to be confounded with supra-annual seasonality, as the STL algorithm is robust to a range of seasonality frequencies (Cleveland et al. 1990). For all species, the algorithm successfully decomposed the raw phenological time series. Moreover, STL enabled us to extract the contribution of inter-annual variation to phenological time series measured at the monthly scale, as in Vogado et al. (2022). Thus, by not reducing inter-annual variation to a single annual value, we were able to retain greater statistical power for our analyses on climate-phenology relationships.

### Trends in climatic variables

Trends in temperature, precipitation and solar irradiance differed based on seasons. Overall, the wet season was found to show a warming trend while the pre-monsoon warm season from March to May became cloudier. Rainfall did not show a declining trend in any of the seasons. Our finding is in line with Jain et al. (2012) which did not find significant trends in southwest monsoon rainfall from Northeast India over a period of several decades, even though Goswami & Prasad (2023) found a decline in some districts of Arunachal Pradesh bordering the study area.

Equivocal findings on rainfall trends from the region (e.g. Dash et al. 2012, Choudhury et al. 2019) could reflect the importance of topographic diversity (Houze 2012) and landuse (Pielke Sr. et al. 2007) in shaping spatio-temporally fine scale precipitation patterns. Our findings also emphasize the need for taking seasonal variation in long-term climatic patterns into consideration (e.g. as in Jain et al. 2013 and Wagholikar et al. 2014).

### Effects of ENSO and climate on inter-annual variation in tree reproductive phenology

MEI had a strong positive effect on flowering, in accordance with our expectations. The effect of the Multivariate ENSO Index was found to be positive and statistically significant, with flowering responding to periods of El Niño without any lags in response. Similar positive responses of flowering to El Niño have been observed at a number of tropical forest sites (Wright & Calderón 2006, Sakai et al. 2006, Detto et al. 2018).

However, we did not find that the increase in flowering during El Niño was mediated by an increase in solar irradiance, or even any other climate variable. Solar irradiance was actually found to decrease, with rainfall increasing, during periods of El Niño. In general, however, phases of El Niño during the study period did not appear to coincide with significant fluctuations in temperature, precipitation and solar irradiance.

Spatio-temporal variability in ENSO effects on wet season rainfall is widely acknowledged (Kumar et al. 2006). Kumar & Singh (2021) found increased monsoon rainfall over periods of El Niño over Northeast India. Spatio-temporal variability in ENSO-monsoon rainfall relationships is likely to be widespread over the Indian subcontinent (Hrudya et al. 2020). ENSO-climate coupling in the Indian Ocean region is much more complex than over the Neotropics or Southeast Asia, possibly due to the involvement of the Indian Ocean Dipole (IOD) (Ashok et al. 2001, Ummenhofer et al. 2011) as well as strong local climatic forcing. We did not model the interactive effects of ENSO and IOD on reproductive phenology, primarily to ensure model simplicity and ease of interpretation.

As we had hypothesized, fruiting was found to be positively associated with solar irradiance, albeit exhibiting a general lagged response. The importance of solar irradiance in determining both seasonal and inter-annual patterns underscore its importance in fruit formation and development. We found the proximal cause of both seasonal (Datta et al. 2025) and inter-annual variation in fruiting to be the same (i.e. increased solar irradiance), which is expected in seasonal tropical forests (Wright & Calderón 2006). A number of studies have found solar irradiance to have the strongest effect on inter-annual fruiting intensity (Wright & Calderón 2006, Chapman et al. 2018). ENSO effects were found to be negligible, unlike in the Neotropics (Wright et al. 1999, Wright & Calderón 2006). Interestingly, Chapman et al. (2018) found fruiting in Kibale (Uganda) to increase during periods of El Niño, and also in relation to increases in solar irradiance, but there was no significant effect of ENSO on solar irradiance itself. The longer term pattern in Kibale is that of increasing rainfall during El Niño episodes (Chapman et al. 2018) which is similar to what we found. This is contrary to that observed in the Neotropics or in Southeast Asia (Ropelewski & Halpert 1987, which also highlighted ENSO-based droughts in India, Aceituno 1988). Thus, the absence of the El Niño-solar irradiance-plant reproduction mechanistic pathway hypothesized by Wright & Calderón (2006) is because El Niño episodes are not associated with increases in solar irradiance in our site.

The strong relationship between ENSO and flowering intensity in the absence of mediating effects by climate variables is harder to explain. Inter-annual variation in flowering intensity appeared to be independent of temperature, precipitation and solar irradiance. However, it is possible that flowering was responding to finer-scale variation in climatic conditions, which we could not test as our phenological data was analyzed on a monthly scale. Community-level flowering was much more synchronous than fruiting in the study area, which can be inferred from the lower dispersion of flowering time as indicated by circular statistical measures (Datta et al. 2025). This means that inter-annual variation in flowering can possibly be triggered by climatic fluctuations occurring over a much smaller scale, such as a few days of reduced temperature or precipitation anomalies.

In addition, flowering patterns at the seasonal scale are closely associated with those of leaf flushing in the site (Datta et al. 2025), as well as in drier tropical forests in general (Murali & Sukumar 1994, Kushwaha et al. 2011). Daylength appears to be the principal proximate cue for leaf flushing in the site (Datta et al. 2025), although severe reductions in water availability in drought years could cause leaf flushing intensity to decrease. We did not observe any such patterns, possibly because water availability was not a limiting factor in the study area, owing to its location in the Himalayan foothills, where the water table is known to be high.

### Traits mediating species-specific responses in phenological intensity to ENSO/climate

Contrary to our expectations, fruit length or width was not found to affect species-specific responses to inter-annual variation in solar irradiance. We also did not find the fruiting intensity of mechanically-dispersed species to be more sensitive to changes in solar irradiance. However, species with a fruiting peak in October appeared to show a greater sensitivity to solar irradiance. There were five species (viz. *Beilschmiedia assamica*, *L. jenkinsiana*, *M. hodgsonii*, *S. tetragonum* and *Dalrympelea pomifera*) whose peak fruiting was in October. The increase in solar irradiance in this month, after the withdrawal of the southwest monsoon, is likely to be responsible for the secondary peak in the bimodal distribution of seasonal fruiting in the site (Datta et al. 2025). Furthermore, solar irradiance may represent a limiting resource for species that fruit during this period. The months from November to February consist of cool, dry conditions with reduced daylength. This is a period of low phenological activity (Datta et al. 2025). Therefore, species that fruit in the post-monsoon period can be expected to show greater sensitivity to fluctuations in solar irradiance. Species that fruit during the warm dry season months of March to May did not show an increased sensitivity to inter-annual variation in solar irradiance. These months coincide with the seasonal peak in fruiting (Datta et al. 2025), and have shown a declining trend in solar irradiance. This is possibly because solar irradiance is not a limiting resource at this time of the year. The lack of morphological or reproductive trait-specific long-term phenological responses to climatic variability could indicate that flower/fruit sizes are less of a constraint in reproductive activity than solar irradiance itself.

### A holistic view of complex seasonal and inter-annual phenological patterns

Phenological patterns in the tropics are highly complex, and we emphasize the need to view flowering and fruiting timeseries as combinations of cyclic patterns with varying frequencies and amplitudes. Contrary to long-held beliefs arising out of the lack of a defined growing period, seasonality in plant phenology is widespread in the tropics (van Schaik et al. 1993, Morellato et al. 2000, Perina et al. 2019), including in the Eastern Himalaya (Datta et al. 2025). In this study, we explicitly controlled for seasonality using time series decomposition or de-seasonalized residuals, enabling inter-annual variation in flowering and fruiting to be distinguished from seasonal patterns. The dominance of nonlinear patterns over relatively monotonic trends could indicate that inter-annual variations in phenology are closely associated with similar fluctuations in climate and that prevailing climatic extremes are not yet causing steep community-wide declines in phenological activity. Nevertheless, climatic extremes, whose prevalence is increasing due to climate change (Hennessy et al. 1997, Guhathakurta et al. 2011, Yaduvanshi et al. 2021) can cause sudden declines (Butt et al. 2015). Climatic fluctuations can also disrupt the usual patterns of phenological seasonality and cyclicity (Chapman et al. 2018). In Pakke, fluctuations in flowering have been more marked in recent years, with the long-term lows in 2022 and 2024 and a year of very high flowering in 2023. Wright et al. (1999) found years of very low fruiting following El Niño-induced peaks in Panama, possibly owing to reduced energy availability after large reproductive investment in the previous year. However, in Pakke, these flowering lows coincided with La Niña events, thus confounding effects of climate-phenology coupling and trade-offs pertaining to energetic investment. Discerning the imprint of climate change upon trends in tropical phenology can be difficult, given the intrinsic complexity in tropical phenological patterns and the wide variation in species-specific responses.

## CONCLUSION

We feel that our study is a valuable addition to the growing body of work on long-term patterns of flowering and fruiting intensity in the tropics. Although the importance of studying long-term flowering and fruiting patterns in the tropics is gaining increasing importance, our study is among the first to explicitly model the effect of climatic variability on inter-annual patterns of flowering and fruiting intensity using a mechanistic framework that examined whether climatic variables mediate the effects of ENSO on reproductive phenological intensity. This is also among the first studies to explicitly incorporate traits in modelling species-specific inter-annual phenological responses. Future research should aim for more mechanistic examinations of long-term climate-phenology relationships, with a focus on trade-offs in energetic investment. Such approaches are still highly lacking in tropical phenological research.

## Supporting information

Supplementary Files_Banerjee&Datta

## ACKNOWLEDGEMENTS

We express our gratitude to the Arunachal Pradesh Forest Department for providing research permits and for their overall support, especially Pekyom Ringu, G.N. Sinha and Bharat Bhatt for their support in the past and N.Tam, Dr. Damodhar A.T., Millo Tasser and Tajum Yomcha for their continuing support. We are grateful to the Pakke Tiger Reserve management (in particular, Tana Tapi, Satyaprakash Singh, Dhawan Kumar Rawat, P.B. Rana, Rubu Tado, Kime Rambia and many frontline staff of the reserve) for facilitating this work over many years. This work would not have been possible without the contributions of the field technicians and assistants of the Eastern Himalaya Programme, Nature Conservation Foundation over the years. These include late Kumar Thapa, late Tali Nabum, late Narayan Mogar, Khem Thapa, Turuk Brah, Peter Wage, Jacob Brah, Sagar Kino, Arjun Rai, and Sital Dako. Current and former researchers of EHP-NCF who played a key role in the tree phenology and climate data collection include Bibidishananda Basu, Noopur Borawake, Dr. Rohit Naniwadekar, Dr. Akanksha Rathore, Swati Sidhu, Amruta Rane, Ushma Shukla, Devathi Parashuram, and Saniya Chaplod among others. We thank Rom Whitaker and the Agumbe Rainforest Research Station for donating the weather station in 2011 and Shankar CM for repeated technical support. We are grateful to Kamal Sonar for maintaining the weather station for several years. Arjun Rai, Sital Dako, Veena Rai and Chaithra Gowda helped with data entry. Grants that helped us carry out our long-term research and conservation programme in the Eastern Himalaya include the National Geographic Society, Disney Wildlife Conservation Fund, Wildlife Conservation Society, Whitley Fund for Nature, the Serenity Trust and the British Ecological Society grant.

## AUTHOR CONTRIBUTIONS

AD conceived, initiated and acquired funding for the project. SB and AD participated in field data collection along with the EHP-NCF field team. Data curation was done by AD and SB. SB completed formal data analyses with contributions from AD. SB and AD wrote and reviewed the paper.

## DATA AVAILABILITY STATEMENT

The raw phenology and climate data has been archived at Zenodo and can be accessed using the following DOI: https://doi.org/10.5281/zenodo.18336408. R code used forstatistical analyses will be archived with this submission prior to publication.

