## Supplementary Files_Banerjee&Datta for "Climate-linked divergence in tree flowering and fruiting in an Eastern Himalayan tropical forest"

| Biological level of phenological trend assessment | GAM model outputs | |
| --- | --- | --- |
| **Flowering at community level or seed dispersal modes** | 2011-2020 | 2021-2024 |
| Community level | edf=3.424, F=3.986, p=0.005* | edf=6.951, F=11.86, p<0.0001* |
| Bird-dispersed species | edf=3.19, F=3.032, p=0.022* | edf=4.671, F=3.401, p=0.01* |
| Mammal-dispersed species | edf=2.563, F=3.451, p=0.017* | edf=8.715, F=14.66, p<0.0001* |
| Mechanically-dispersed species | edf=9.185, F=3.955, p=0.0001* | edf=1.784, F=1.531, p=0.231 |
| **Species-level flowering** |  |  |
| *Actinodaphne obovata* | edf=9.744, F=4.297, p<0.0001* | edf=8.776, F=17.36, p<0.0001* |
| *Aglaia spectabilis* | edf=1, F=0.004, p=0.949 | edf=7.907, F=9.299, p<0.0001* |
| *Ailanthus integrifolia* | edf=1.001, F=1.136, p=0.289 | edf=8.662, F=20.65, p<0.0001* |
| *Artocarpus chama* | edf=14.12, F=7.599, p<0.0001* | edf=8.492, F=15.57, p<0.0001* |
| *Alstonia scholaris* | edf=12.75, F=5.889, p<0.0001* | edf=8.734, F=19.32, p<0.0001* |
| *Baccaurea ramiflora* | edf=13.43, F=5.626, p<0.0001* | edf=4.422, F=2.971, p=0.027* |
| *Bauhinia variegata* | edf=3.216, F=1.693, p=0.158 | edf=6.718, F=6.126, p=0.002* |
| *Beilschmiedia assamica* | edf=1.001, F=1.726, p=0.192 | edf=5.095, F=8.148, p<0.0001* |
| *Beilschmiedia sp* | edf=2.711, F=3.892, p=0.009* | edf=7.397, F=9.383, p<0.0001* |
| *Canarium resiniferum* | edf=1, F=1.455, p=0.231 | edf=8.006, F=18.03, p<0.0001* |
| *Chisocheton cumingianus* | edf=2.168, F=1.408, p=0.247 | edf=3.008, F-2.734, p=0.043* |
| *Choerospondias axillaris* | edf=3.272, F=2.348, p=0.059 | edf=2.165, F=1.76, p=0.121 |
| *Chukrasia tabularis* | edf=3.369, F=3.37, p=0.011* | edf=1, F=0.669, p=0.418 |
| *Cryptocarya sp.* | edf=2.548, F=0.937, p=0.44 | edf=1.149, F=2.236, p=0.115 |
| *Dalrympelea pomifera* | edf=13.43, F=5.055, p<0.0001* | edf=8.749, F=18.15, p<0.0001* |
| *Dillenia indica* | edf=1.097, F=1.464, p=0.209 | edf=1.305, F=4.25, p=0.056 |
| *Epicharis cuneata* | edf=6.845, F=3.635, p<0.0001* | edf=7.75, F=12.41, p<0.0001* |
| *Dysoxylum gotadhora* | edf=14.62, F=7.376, p=0.053 | edf=8.01, F=5.723, p=0.0001* |
| *Prasoxylon excelsum* | edf=7.963, F=8.206, p<0.0001* | edf=7.705, F=8.118, p<0.0001* |
| *Endospermum chinense* | edf=1, F=4.932, p=0.029* | edf=7.800, F=12.18, p<0.0001* |
| *Gmelina arborea* | edf=1.319, F=3.174, p=0.060 | edf=1, F=1.685, p=0.202 |
| *Gynocardia odorata* | edf=1.835, F=2.066, p=0.122 | edf=8.699, F=15.05, p<0.0001* |
| *Horsfieldia kingii* | edf=1.002, F=6.163, p=0.015* | edf=5.017, F=3.736, p=0.009* |
| *Knema angustifolia* | edf=7.644, F=5.519, p<0.0001* | edf=8.691, F=15.5, p<0.0001* |
| *Livistona jenkinsiana* | edf=14.65, F=6.364, p<0.0001* | edf=8.804, F=17.64, p<0.0001* |
| *Magnolia hodgsonii* | edf=14.25, F=7.013, p<0.0001* | edf=8.854, F=22.59, p<0.0001* |
| *Monoon simiarum* | edf=13.7, F=5.355, p<0.0001* | edf=8.859, F=30.52, p<0.0001* |
| *Picrasma javanica* | edf=2.289, F=2.07, p=0.12 | edf=8.423, F=13.08, p<0.0001* |
| *Pterospermum acerifolium* | edf=7.864, F=5.156, p<0.0001* | edf=8.613, F=13.73, p<0.0001* |
| *Pterygota alata* | edf=18.16, F=13.35, p<0.0001* | edf=2.385, F=2.129, p=0.109 |
| *Prunus ceylanica* | edf=1, F=0.659, p=0.419 | edf=8.636, F=15.02, p<0.0001* |
| *Sterculia villosa* | edf=2.07, F=2.047, p=0.11 | edf=1.661, F=0.318, p=0.691 |
| *Stereospermum tetragonum* | edf=14.79, F=6.253, p<0.0001* | edf=8.369, F=18.24, p<0.0001* |
| *Tetrameles nudiflora* | edf=3.341, F=1.728, p=0.157 | edf=8.548, F=12.13, p<0.0001* |
| *Zanthoxylum rhetsa* | edf=12.07, F=4.228, p<0.0001* | edf=7.629, F=8.87, p<0.0001* |
| **Fruiting at community level or seed dispersal modes** |  |  |
| Community level | edf=1.925, F=0.724, p=0.415 | edf=8.902, F=37.28, p<0.0001* |
| Bird-dispersed species | edf=1.689, F=3.634, p=0.03* | edf=8.895, F=35.68, p<0.0001* |
| Mammal-dispersed species | edf=2.144, F=0.826, p=0.365 | edf=8.865, F=29.23, p<0.0001* |
| Mechanically-dispersed species | edf=14.67, F=7.503, p<0.0001* | edf=1, F=13.27, p=0.001* |
| **Species level** |  |  |
| *Actinodaphne obovata* | edf=1, F=1.733, p=0.191 | edf=8.821, F=24.29, p<0.0001* |
| *Aglaia spectabilis* | edf=7.907, F=9.299, p<0.0001* | edf=2.307, F=3.302, p=0.035* |
| *Ailanthus integrifolia* | edf=14.480, F=5.655, p<0.0001* | edf=7.784, F=13.67, p<0.0001* |
| *Artocarpus chama* | edf=18.420, F=20.49, p<0.0001* | edf=8.763, F=18.42, p<0.0001* |
| *Alstonia scholaris* | edf=2.449, F=2.779, p=0.04* | edf=7.922, F=20.78, p<0.0001* |
| *Baccaurea ramiflora* | edf=4.666, F=2.787, p=0.021* | edf=1, F=0.4, p=0.531 |
| *Bauhinia variegata* | edf=14, F=5.947,p<0.0001* | edf=7.990, F=20.16, p<0.0001* |
| *Beilschmiedia assamica* | edf=1, F=0.478, p=0.49 | edf=8.813, F=19.78, p<0.0001* |
| *Beilschmiedia sp* | edf=14.57, F=7.134, p<0.0001* | edf=1, F=7.148, p=0.012* |
| *Canarium resiniferum* | edf=1.002, F=1.699, p=0.195 | edf=8.893, F=35.47, p<0.0001* |
| *Chisocheton cumingianus* | edf=1.001, F=4.239, p=0.043* | edf=6.825, F=7.106, p<0.0001* |
| *Choerospondias axillaris* | edf=4.815, F=3.902, p=0.002* | edf=8.781, F=21.86, p<0.0001* |
| *Chukrasia tabularis* | edf=3.996, F=2.039, p=0.088 | edf=8.850, F=26.59, p<0.0001 |
| *Cryptocarya sp.* | edf=2.510, F=1.726, p=0.176 | edf=8.500, F=24.99, p<0.0001* |
| *Dalrympelea pomifera* | edf=14.17, F=8.147, p<0.0001* | edf=8.771, F=17.67, p<0.0001* |
| *Dillenia indica* | edf=3.441, F=2.798. p=0.028* | edf=8.670, F=24.23, p<0.0001* |
| *Epicharis cuneata* | edf=1, F=0.059, p=0.808 | edf=8.780, F=25.41, p<0.0001 |
| *Dysoxylum gotadhora* | edf=1.001, F=1.771, p=0.186 | edf=8.869, F=26.45, p<0.0001* |
| *Prasoxylon excelsum* | edf=5.330, F=1.789, p=0.104 | edf=7.911, F=30.54, p<0.0001* |
| *Endospermum chinense* | edf=1.001, F=1.846, p=0.178 | edf=1, F=9.425, p=0.004* |
| *Ficus drupacea* | edf=1.984, F=1.079, p=0.322 | edf=8.852, F=21.84, p<0.0001* |
| *Gmelina arborea* | edf=14.700, F=13.09, p<0.0001* | edf=3.494, F=7.825, p=0.0002* |
| *Gynocardia odorata* | edf=2.691, F=3.154, p=0.026* | edf=8.831, F=19.27, p<0.0001* |
| *Horsfieldia kingii* | edf=3.374, F=1.876, p=0.119 | edf=8.700, F=33.16, p<0.0001* |
| *Knema angustifolia* | edf=6.223, F=5.664, p<0.0001* | edf=8.78, F=23.14, p<0.0001* |
| *Livistona jenkinsiana* | edf=14.54, F=8.61, p<0.0001* | edf=8.744, F=15.66, p<0.0001* |
| *Magnolia hodgsonii* | edf=4.638, F=2.174, p=0.056 | edf=8.260, F=24.24, p<0.0001* |
| *Monoon simiarum* | edf=2.427, F=3.128, p=0.030* | edf=1, F=1.34, p=0.254 |
| *Picrasma javanica* | edf=13.870, F=5.752, p<0.0001* | edf=8.692, F=16.84, p<0.0001* |
| *Pterospermum acerifolium* | edf=4.142, F=3.339, p=0.008* | edf=5.558, F=3.606, p=0.01* |
| *Pterygota alata* | edf=12.160, F=4.604, p<0.0001* | edf=8.159, F=8.457, p<0.0001* |
| *Prunus ceylanica* | edf=2.497, F=3.212, p=0.025* | edf=3.127, F=4.733, p=0.005* |
| *Sterculia villosa* | edf=6.501, F=3.015, p=0.005* | edf=8.845, F=24.28, p<0.0001* |
| *Stereospermum tetragonum* | edf=1.476, F=6.288, p<0.05* | edf=5.550, F=6.923, p<0.05* |
| *Tetrameles nudiflora* | edf=2.193, F=1.138, p=0.35 | edf=4.120, F=5.398, p=0.001* |
| *Zanthoxylum rhetsa* | edf=1.002, F=10.06, p=0.002* | edf=8.407, F=15.62, p<0.0001* |

Table S1: Model outputs from cubic smoothed splines fit to de-seasonalized time series of flowering and fruiting intensity using Generalized Additive Modelling (GAM). Edf represents the effective degrees of freedom and the significance of the F statistic is tested against a null hypothesis of the smooth term having no effect on the response. Smoothed splines were fit to flowering and fruiting data for each of 36 species for which at least 10 trees were monitored and also at the community level and for each of the principal seed dispersal modes (viz. Bird-dispersed, Mammal-dispersed and Mechanically-dispersed species).

| Variable | Beta coefficient (95% CI) of trend component | | |
| --- | --- | --- | --- |
|  | Warm dry | Wet | Cool dry |
| Mean maximum temperature | -0.204 (-0.472, 0.064) | 0.191 (0.040, 0.343) | -0.184 (-0.391, 0.022) |
| Mean minimum temperature | -0.061 (-0.506, 0.384) | 0.080 (0.051, 0.109) | -0.135 (-0.335, 0.066) |
| Total monthly rainfall | 1.354 (-0.279, 2.987) | 0.847 (-0.928, 2.623) | 0.218 (-0.079, 0.515) |
| Mean maximum solar irradiance | -21.718 (-37.725, -5.711) | -8.601 (-19.673, 2.471) | -2.096 (-7.569, 3.377) |
| Monthly proportion of rainy days | 0.017 (-0.022, 0.056) | 0.010 (-0.002, 0.023) | 0.001 (-0.013, 0.016) |

Table S2: Model outputs from linear models representing the trend of change in seasonal mean values of climate variables between January 2011 and December 2024 in Pakke Tiger Reserve, Arunachal Pradesh, India.

| Model | AICc | delAIC | AICc weight |
| --- | --- | --- | --- |
| MEI_lag0 | 6546.095 | 0 | 0.9971 |
| MEI_lag1 | 6557.774 | 11.679 | 0.0003 |
| MEI_lag2 | 6573.665 | 27.571 | Negligible |
| MEI_lag6 | 6576.416 | 30.322 | Negligible |
| MEI_lag5 | 6576.615 | 30.520 | Negligible |
| MEI_lag7 | 6576.968 | 30.873 | Negligible |
| MEI_lag3 | 6577.938 | 31.844 | Negligible |
| MEI_lag4 | 6578.849 | 32.754 | Negligible |

Table S3: Model comparison table for MEI-flowering relationships using the Akaike Information Criterion corrected for small sample size (AICc). Each model refers to the value of the MEI variable which reflects the corresponding lagged response of flowering in months, i.e. MEI_lag1: value of MEI in any month that corresponds to a 1-month lagged response in flowering. Thus, the variable MEI_lag1 (which was the only predictor in the model of the same name) had the value of MEI in January 2011 when the value of the flowering response variable was that of February 2011, and so on.

| Model | AICc | delAICc | AICc weight |
| --- | --- | --- | --- |
| meanmax_lag0 | 6578.648 | 0 | 0.247 |
| solarRad_lag0 | 6578.764 | 0.116 | 0.233 |
| solarRad_ lag8 | 6579.210 | 0.562 | 0.186 |
| rainydays_lag0 | 6580.045 | 1.397 | 0.123 |
| rainydays_lag8 | 6580.251 | 1.603 | 0.111 |
| totrain_lag8 | 6580.422 | 1.774 | 0.102 |

Table S4: Model comparison table for climate-flowering relationships using the Akaike Information Criterion corrected for small sample size (AICc). Each model refers to the value of the temperature, precipitation or solar irradiance variable which reflects the corresponding lagged response of flowering in months, i.e. solarRad_lag8: value of monthly mean of daily maximum solar irradiance in any month that corresponds to a 8-month lagged response in flowering. Thus, the variable solarRad_lag8 (which was the only predictor in the model of the same name) had the value of solar irradiance in January 2011 when the value of the flowering response variable was that of September 2011, and so on.

| Covariate | Beta coefficient (95% CI) | p value |
| --- | --- | --- |
| Intercept | -3.652 (-4.021, -3.283) | <0.0001* |
| MEI (without lags) | 0.273 (0.176, 0.371) | <0.0001* |
| Flower size | -0.002 (-0.039, 0.036) | 0.087 |
| MEI (without lags)*Flower size | -0.009 (-0.032, 0.014) | 0.293 |
| AR(1) | 1.898 (1.603, 2.193) | <0.0001* |
| AR(12) | 2.932 (2.640, 3.223) | <0.0001* |

| Covariate | Beta coefficient (95%CI) | p value |
| --- | --- | --- |
| Intercept | -3.654 (-4.054, -3.255) | <0.0001* |
| MEI (without lags) | 0.289 (0.157, 0.421) | <0.0001* |
| Bisexual | 0.075 (-0.217, 0.368) | 0.613 |
| Monoecious | -0.384 (-0.883, 0.115) | 0.132 |
| MEI (without lags)*Bisexual | -0.063 (-0.236, 0.109) | 0.473 |
| MEI (without lags)*Monoecious | -0.021 (-0.345, 0.304) | 0.901 |
| AR(1) | 1.885 (1.590, 2.179) | <0.0001* |
| AR(12) | 2.919 (2.628, 3.210) | <0.0001* |

Table S5: Model outputs of GLMMs representing interactive effects between MEI and a) flower size and b) reproduction system (i.e. either Bisexual, Monoecious or Dioecious), on the proportion of trees in flower in Pakke Tiger Reserve, Arunachal Pradesh, India, between 2011 and 2020. Significant effects are indicated with an asterisk (*) and also inferred on the basis of the 95% confidence interval not including zero.

| Model | AICc | delAICc | AICc weight |
| --- | --- | --- | --- |
| solarRad_lag3 | 6078.520 | 0 | 0.791 |
| totrain_lag4 | 6082.903 | 4.38 | 0.088 |
| rainydays_lag0 | 6083.821 | 5.30 | 0.056 |
| solarRad_lag0 | 6084.586 | 6.07 | 0.038 |
| solarRad_lag2 | 6085.261 | 6.74 | 0.027 |

Table S6: Model comparison table for climate-fruiting relationships using the Akaike Information Criterion corrected for small sample size (AICc). Each model refers to the value of the temperature, precipitation or solar irradiance variable which reflects the corresponding lagged response of flowering in months, i.e. solarRad_lag3: value of monthly mean of daily maximum solar irradiance in any month that corresponds to a 3-month lagged response in fruiting. Thus, the variable solarRad_lag3 (which was the only predictor in the model of the same name) had the value of solar irradiance in January 2011 when the value of the flowering response variable was that of April 2011, and so on.

a)

Solar irradiance (3 month lagged response of fruiting)* Sexual reproductive system (i.e. Monoiecy, Dioecy or Bisexuality)

| Covariate | Beta coefficient (95% CI) | p value |
| --- | --- | --- |
| Intercept | -1.597 (-2.846, -0.348) | 0.0128* |
| Solar irradiance (3 month lagged effect) | 0.053 (-0.147, 0.253) | 0.606 |
| Number of trees | 0.021 (-0.039, 0.081) | 0.491 |
| AR(1) | 0.176 (0.125, 0.226) | <0.00018* |
| Bisexual | 0.344 (-0.244, 0.932) | 0.251 |
| Monoecious | 0.460 (-0.452, 1.373) | 0.323 |
| Solar irradiance (3 month lagged effect)* Bisexual | 0.238 (-0.027, 0.503) | 0.079 |
| Solar irradiance (3 month lagged effect)* Monoecious | -0.070 (-0.462, 0.322) | 0.726 |
| SD (Species-specific random intercept) | 0.724 (0.518, 1.014) |  |

b) Solar irradiance (3 month lagged response of fruiting)* Dispersal mode (i.e.Biotic vs Abiotic)

| Covariate | Beta coefficient (95%CI) | p value |
| --- | --- | --- |
| Intercept | -1.368 (-2.502, -0.233) | 0.019* |
| Solar irradiance (3 month lagged response) | 0.130 (0.001, 0.258) | 0.048* |
| Number of trees | 0.021 (-0.038, 0.079) | 0.488 |
| AR(1) | 0.177 (0.127, 0.227) | <0.0001* |
| Abiotic | -0.144 (-0.786, 0.499) | 0.662 |
| Solar irradiance* Abiotic | 0.225 (-0.010, 0.549) | 0.175 |
| SD (Species-specific random intercept) | 0.725 (0.519, 1.012) |  |

c) Solar irradiance (3 month lagged response of fruiting)* Fruit type (i.e. Arillate capsule, Dry pod, Fleshy other, Dry capsule, Drupe)

| Covariate | Beta coefficient (95%CI) | p value |
| --- | --- | --- |
| Intercept | -1.103 (-2.271, 0.064) | 0.064 |
| Solar irradiance (3 month lagged response) | 0.074 (-0.141, 0.288) | 0.500 |
| Number of trees | 0.025 (-0.036, 0.086) | 0.421 |
| AR(1) | 0.176 (0.126, 0.226) | <0.0001* |
| Arillate capsule | -0.363 (-1.120, 0.394) | 0.347 |
| Dry pod | -0.331 (-1.248, 0.585) | 0.479 |
| Fleshy other | -0.668 (-1.370, 0.034) | 0.062 |
| Dry capsule | -0.663 (-1.598, 0.272) | 0.165 |
| Solar irradiance (3 month lagged response)* Arillate capsule | 0.200 (-0.132, 0.532) | 0.238 |
| Solar irradiance (3 month lagged response)* Dry pod | 0.160 (-0.343, 0.664) | 0.533 |
| Solar irradiance (3 month lagged response)* Fleshy other | 0.009 (-0.293, -0.311) | 0.955 |
| Solar irradiance (3 month lagged response)* Dry capsule | 0.367 (-0.077, 0.811) | 0.105 |
| SD (Species-specific random intercept) | 0.684 (0.487, 0.961) |  |

d) Solar irradiance (3 month lagged response of fruiting)*Fruit length

| Covariate | Beta coefficient (95%CI) | p value |
| --- | --- | --- |
| Intercept | -1.216 (-2.399, -0.033) | 0.044 (<0.05)* |
| Solar irradiance (3 month lagged response) | 0.103 (-0.052, 0.258) | 0.194 |
| Number of trees | 0.016 (-0.042, 0.075) | 0.583 |
| AR(1) | 0.173 (0.124, 0.222) | <0.0001* |
| Fruit length | -0.010 (-0.040, 0.021) | 0.529 |
| Solar irradiance (3 month lagged response)* Fruit length | 0.012 (-0.004, 0.029) | 0.150 |
| SD (Species-specific random intercept) | 0.740 (0.530, 1.033) |  |

e) Solar irradiance (3 month lagged response of fruiting)*Fruit width

| Covariate | Beta coefficient (95%CI) | p value |
| --- | --- | --- |
| Intercept | -1.181 (-2.357, -0.005) | 0.049* |
| Solar irradiance (3 month lagged response) | 0.172 (-0.010, 0.354) | 0.064 |
| Number of trees | 0.020 (-0.038, 0.079) | 0.497 |
| AR(1) | 0.173 (0.124, 0.222) | <0.0001* |
| Fruit.width | -0.050 (-0.154, 0.054) | 0.349 |
| Solar irradiance (3 month lagged response)* Fruit width | 0.001 (-0.040, 0.042) | 0.970 |
| SD (Species-specific random intercept) | 0.737 (0.528, 1.037) |  |

Table S7: Model outputs of GLMMs representing interactive effects between solar irradiance and a) reproduction system (i.e. Bisexual, Monoecious or Dioecious) (R^2^: 0.326 (conditional), 0.177 (marginal)), b) Dispersal mode (Biotic vs Abiotic) (R^2^: 0.325 (conditional), 0.173 (marginal)) c) Fruit type (Drupe, Arillate capsule, Fleshy other, Dry pod or Dry capsule (R^2^: 0.330 (conditional), 0.196 (marginal)) d) fruit length (R^2^: 0.332 (conditional), 0.175 (marginal)) e) fruit width (R^2^: 0.326 (conditional), 0.169 (marginal)), on the fruiting intensity in Pakke Tiger Reserve, Arunachal Pradesh, India, between 2011 and 2020. Significant effects are indicated with an asterisk (*) and also inferred on the basis of the 95% confidence interval not including zero.

ENSO effect on solar irradiance: best supported model had 1 month lagged response of solar irradiance to MEI

| Covariate | Beta coefficient (95%CI) | p value |
| --- | --- | --- |
| Intercept | 0.014 (-0.176, 0.205) | 0.881 |
| MEI (1 month lagged response) | -0.200 (-0.402, 0.354) | 0.052 |

ENSO effect on total monthly rainfall: best support model had 1 month lagged response of total monthly rainfall to MEI

| Parameters | Beta (95% CI) | p value |
| --- | --- | --- |
| Intercept | -0.014 (-0.205, 0.177) | 0.888 |
| MEI (1 month lagged response) | 0.190 (-0.012, 0.392) | 0.065 |

Table S8: Outputs of linear model examining the effects of ENSO (as quantified by MEI) on solar irradiance and total rainfall after selecting models representing lagged responses from 0-12 months. The most parsimonious ENSO model for both climate variables represented the 1-month lagged response of solar irradiance and total monthly rainfall to MEI. R^2^ values for the models were 0.027 and 0.023 respectively.

Figure S1

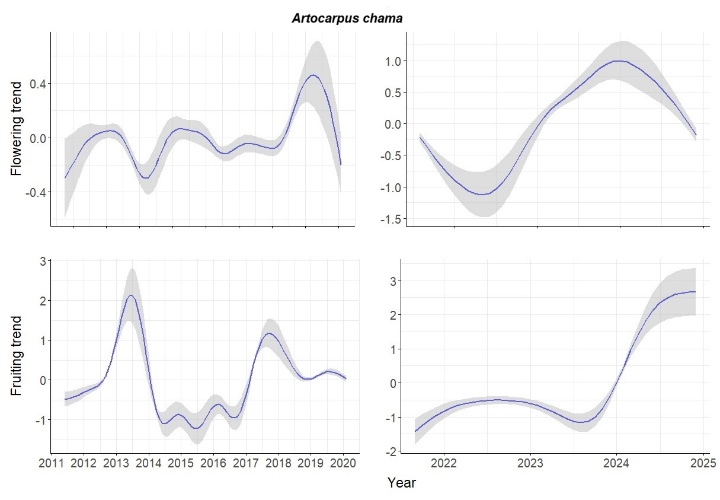

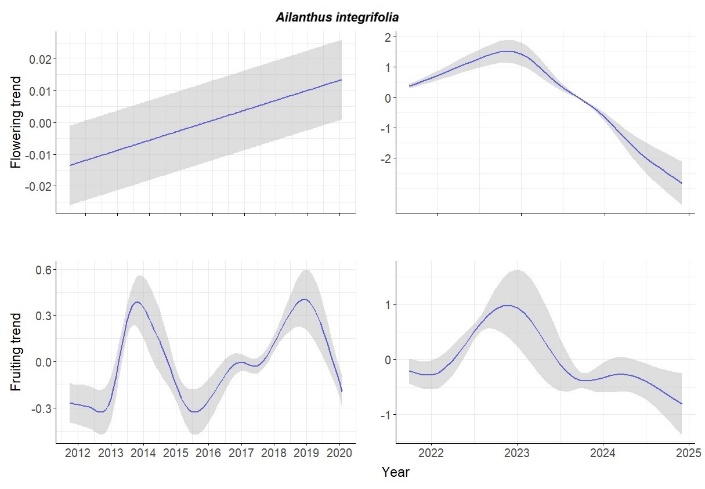

*Artocarpus chama*	*Ailanthus integrifolia*

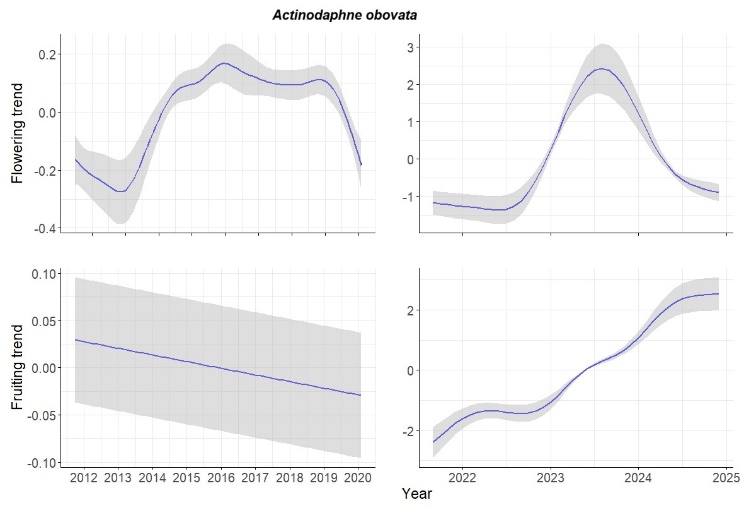

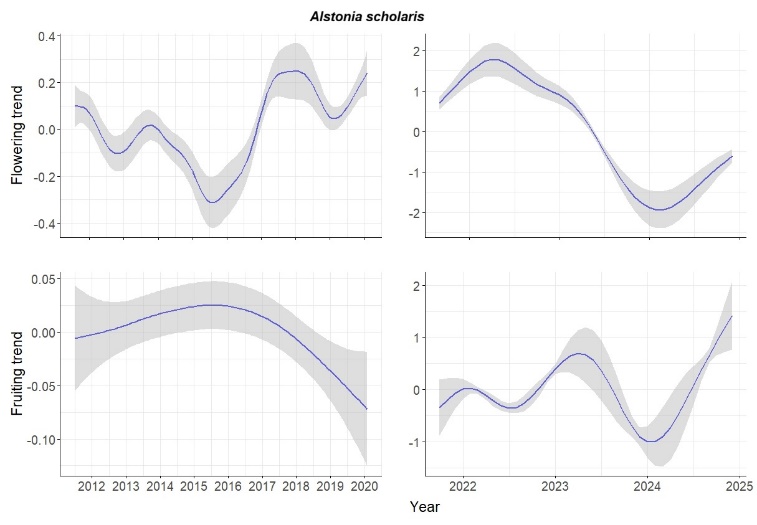

*Actinodaphne obovata*	*Alstonia scholaris*

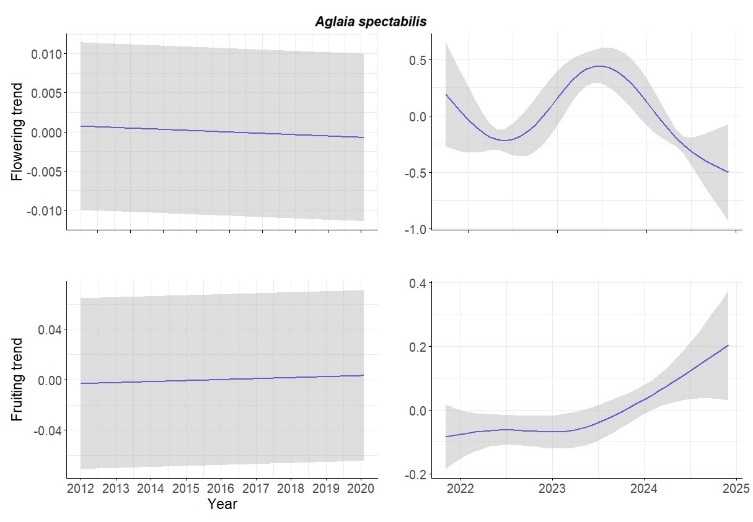

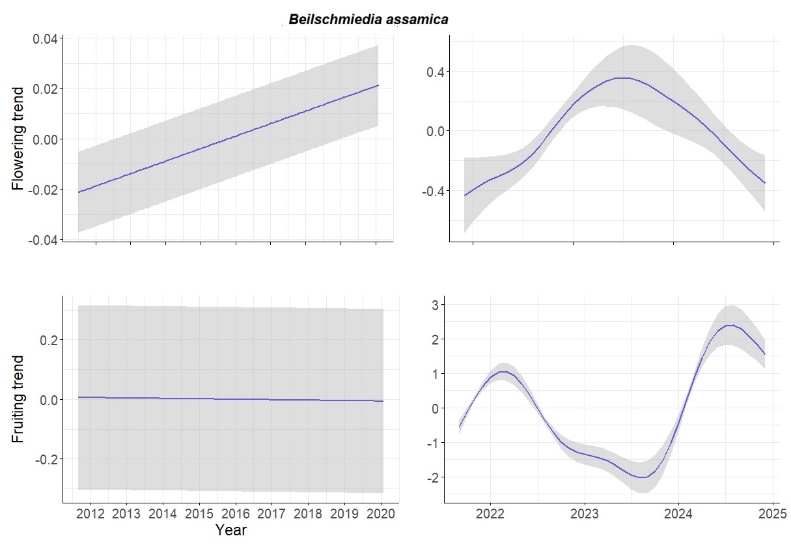

*Aglaia spectabilis	Beilschmiedia assamica*

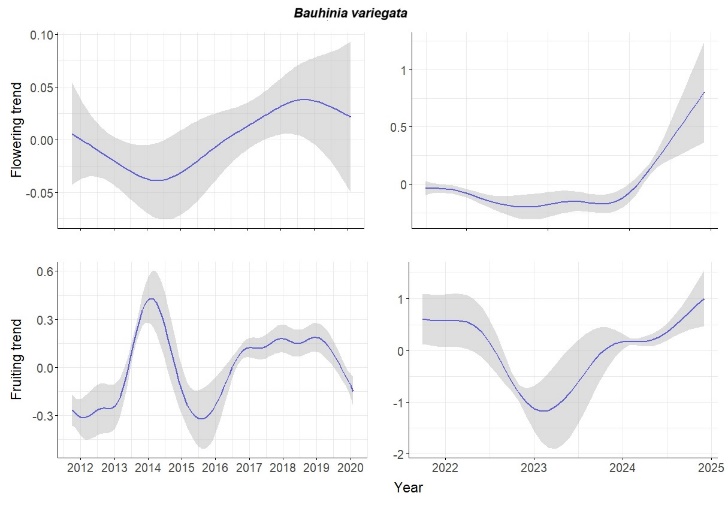

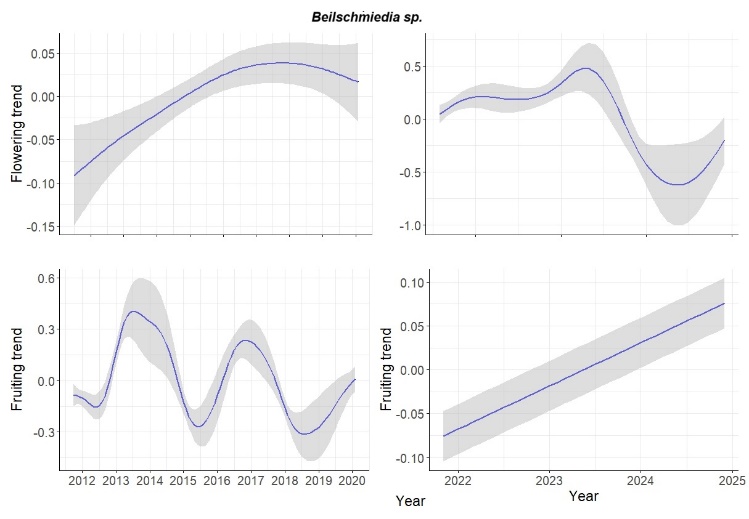

*Bauhinia variegata	Beilschmiedia sp.*

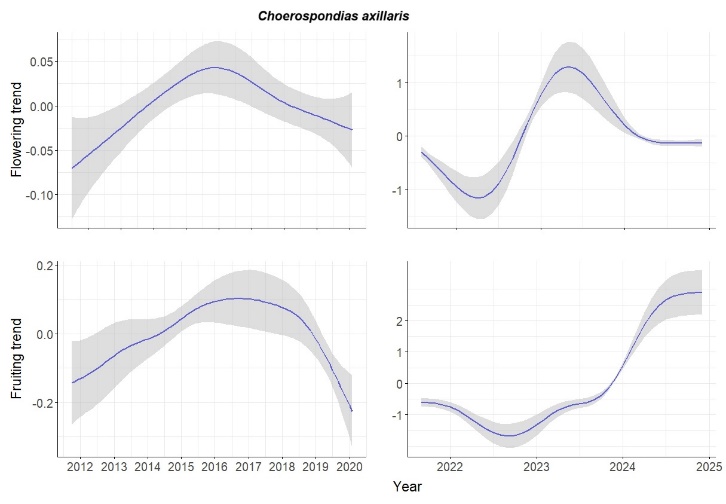

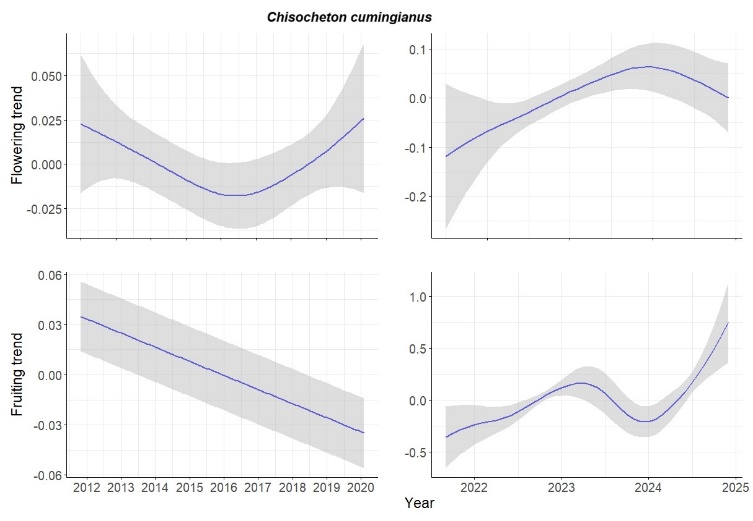

*Choerospondias axillaris	Chisocheton cumingianus*

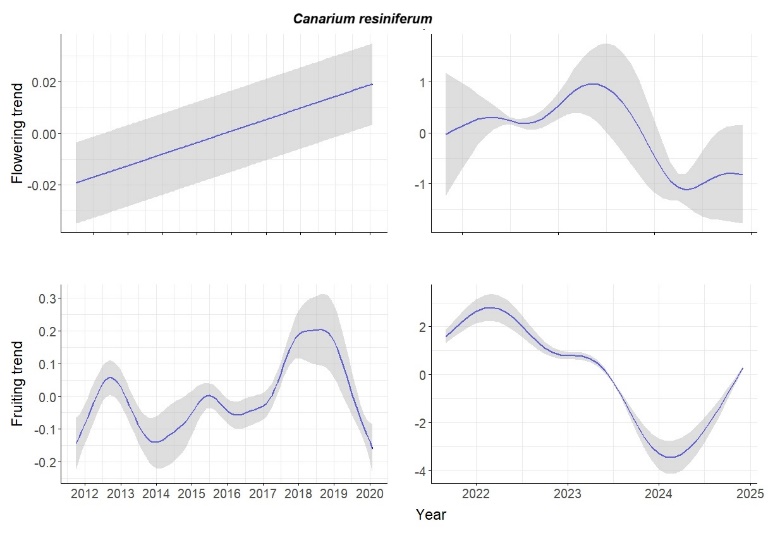

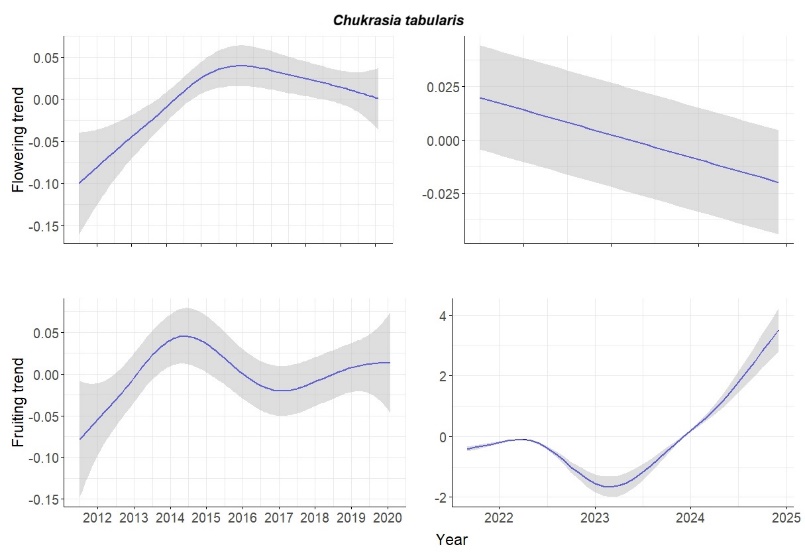
 *Canarium resiniferum	Chukrasia tabularis*

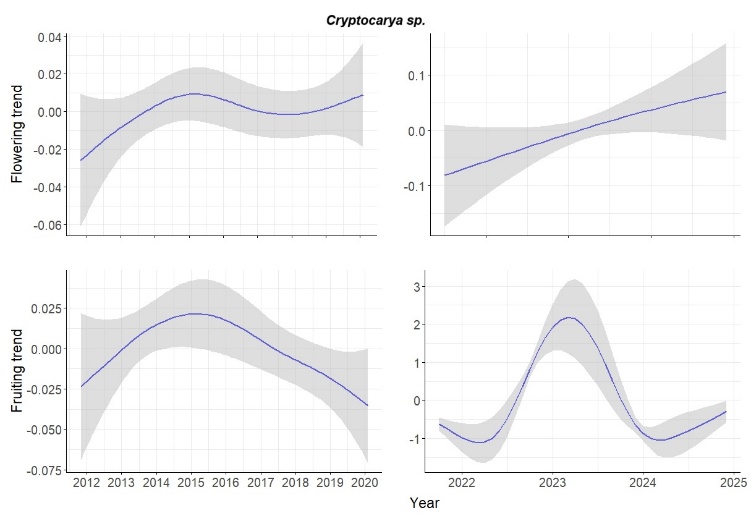

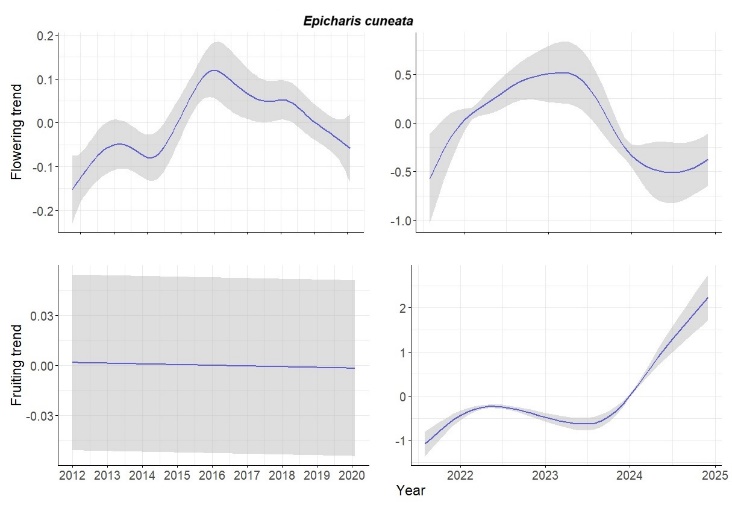

*Cryptocarya sp	Epicharis cuneata*

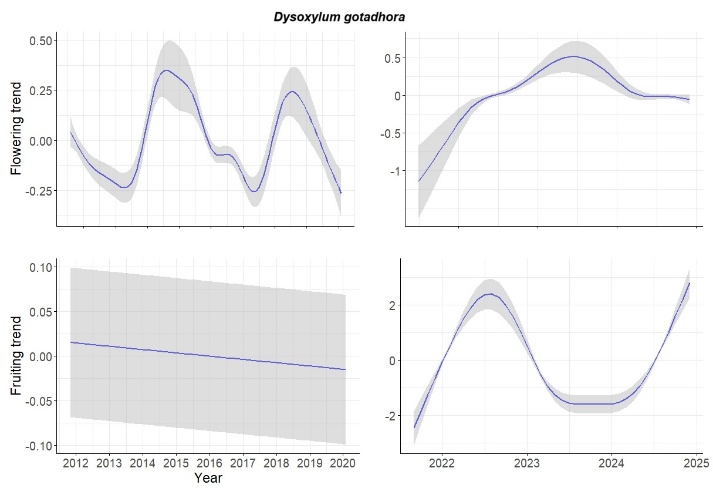

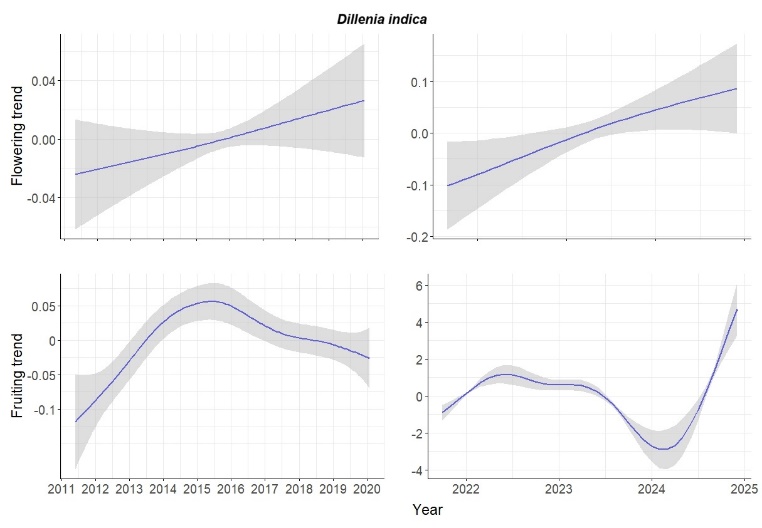

*Dysoxylum gotadhora	Dillenia indica*

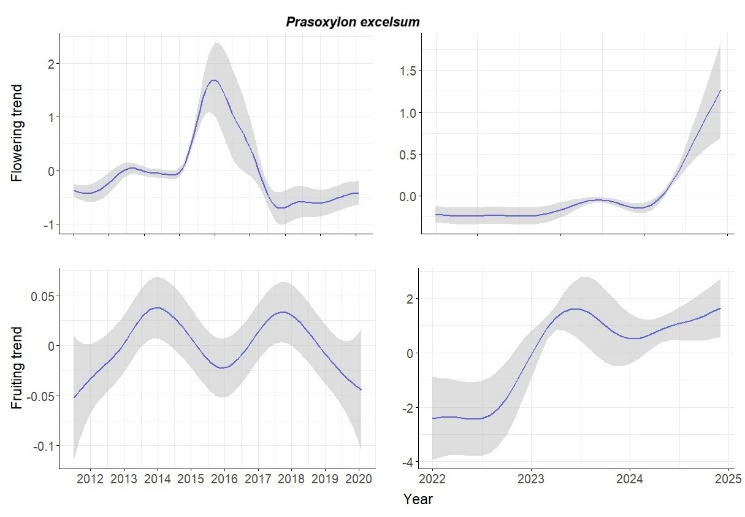

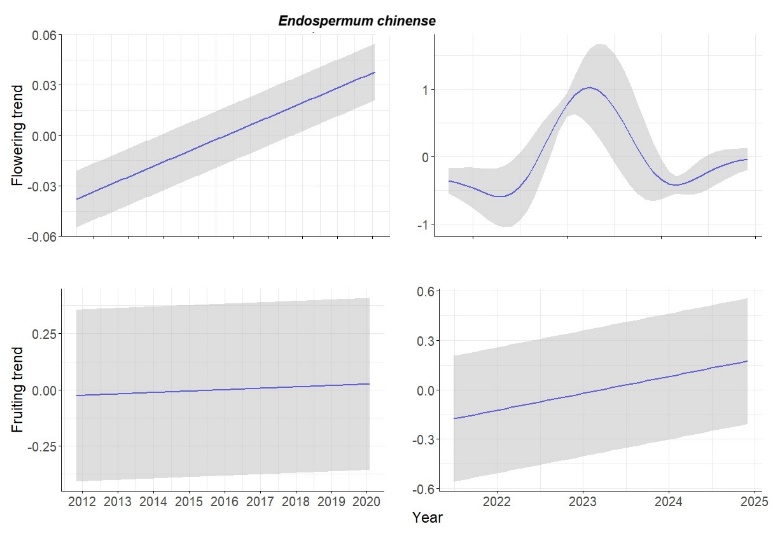

*Prasoxylon excelsum	Endospermum chinense*

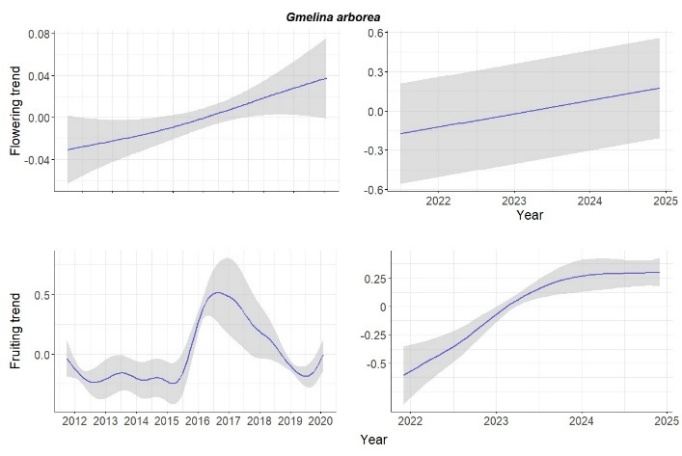

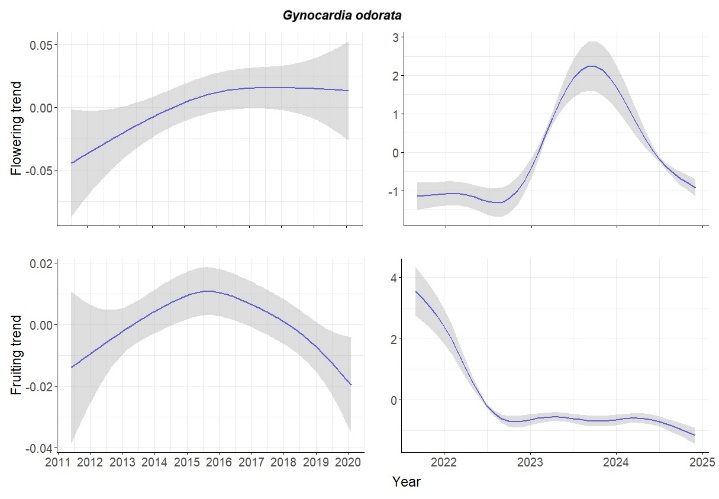

*Gmelina arborea	Gynocardia odorata*

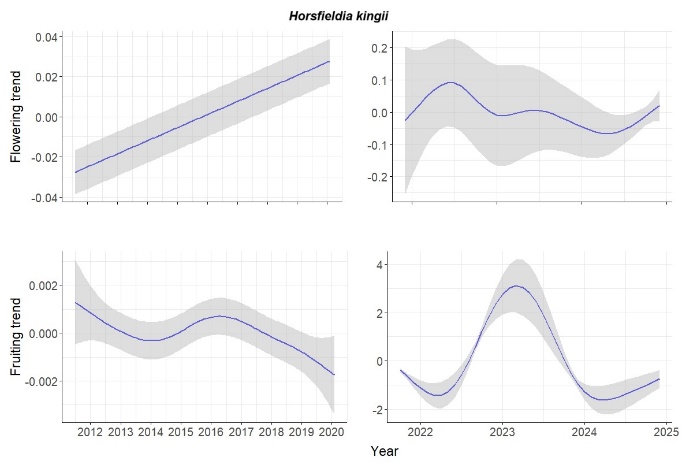

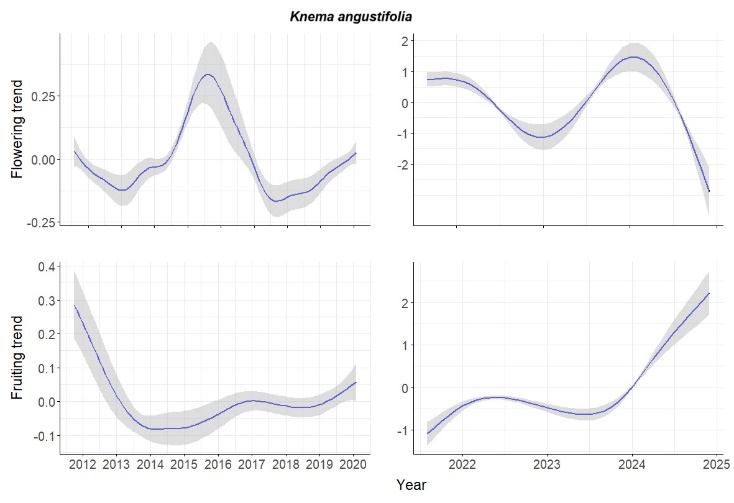

*Horsfieldia kingii	Knema angustifolia*

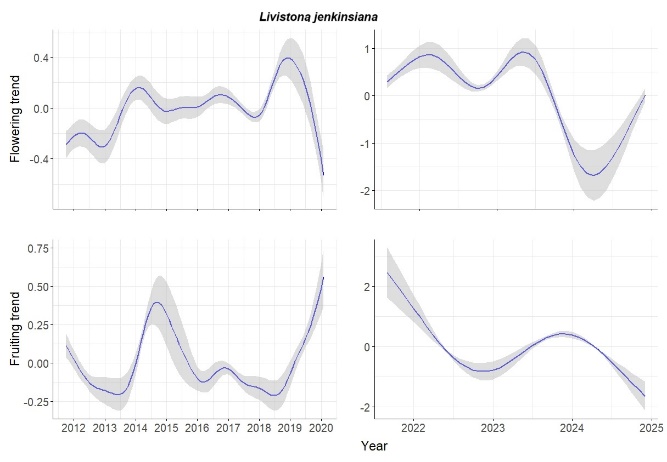

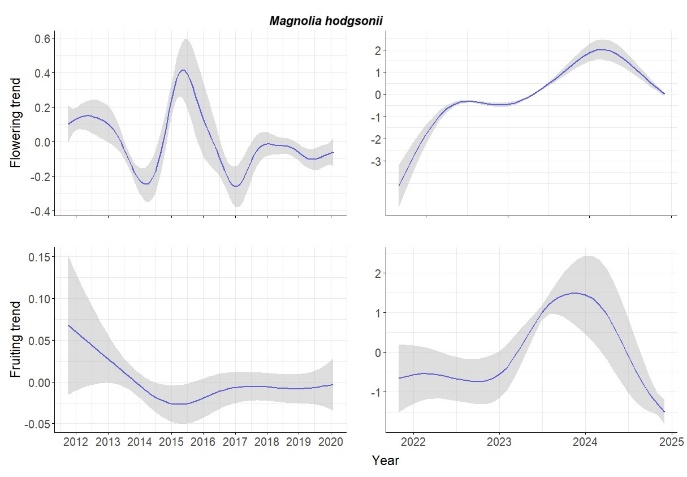

*Livistona jenkinsiana	Magnolia hodgsonii*

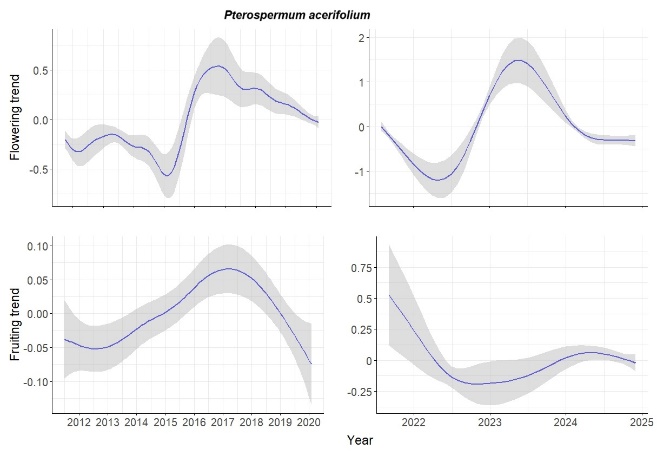

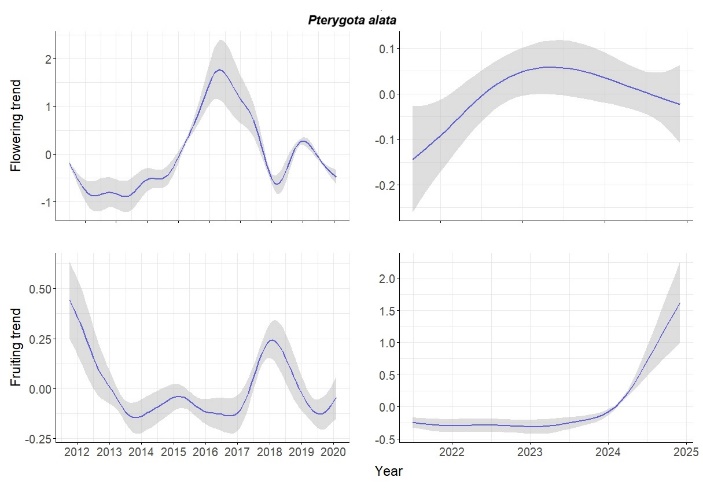

*Pterospermum acerifolium	Pterygota alata*

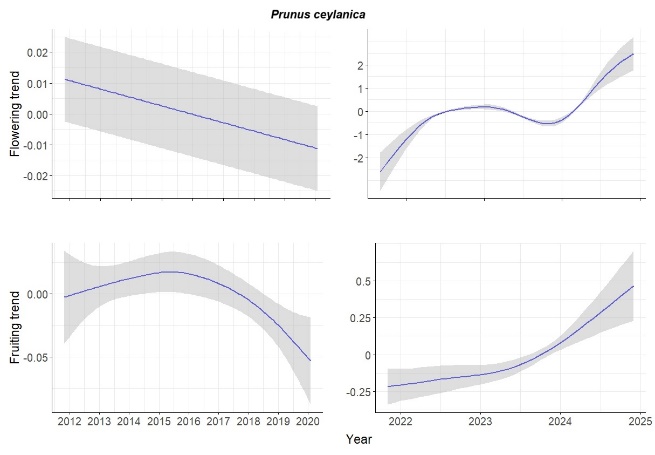

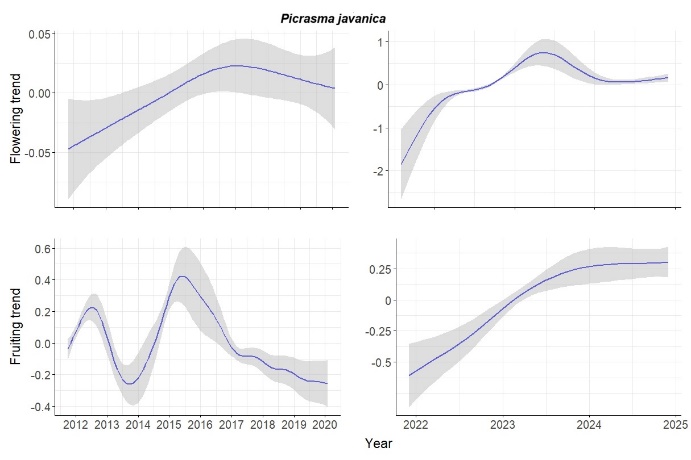

*Prunus ceylanica	Picrasma javanica*

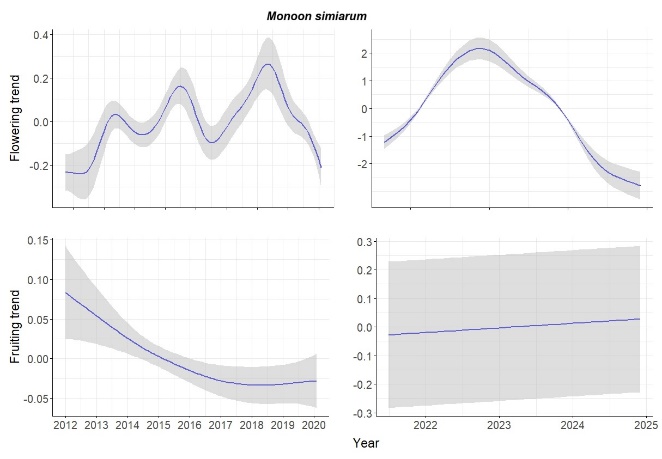

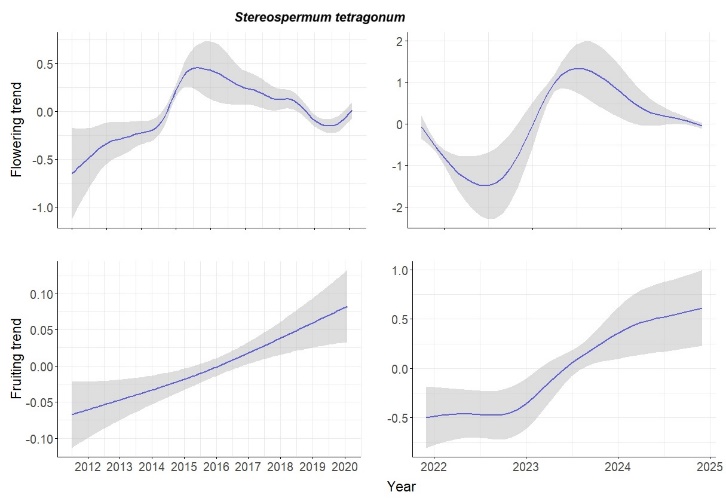

*Monoon simiarum	Stereospermum tetragonum*

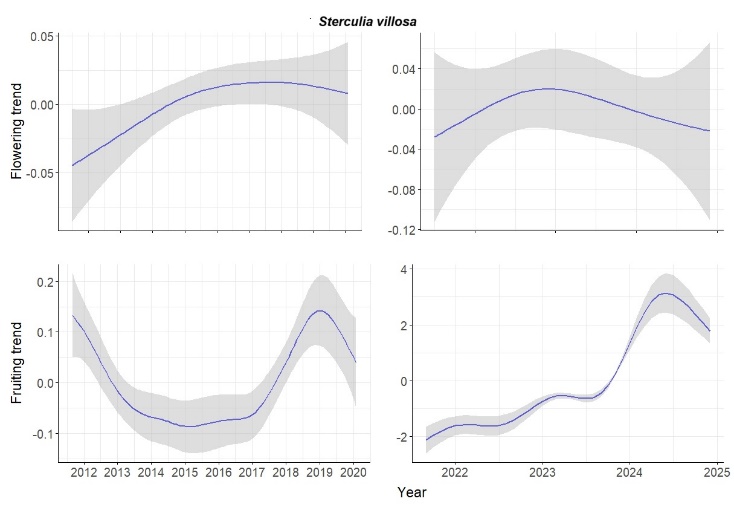

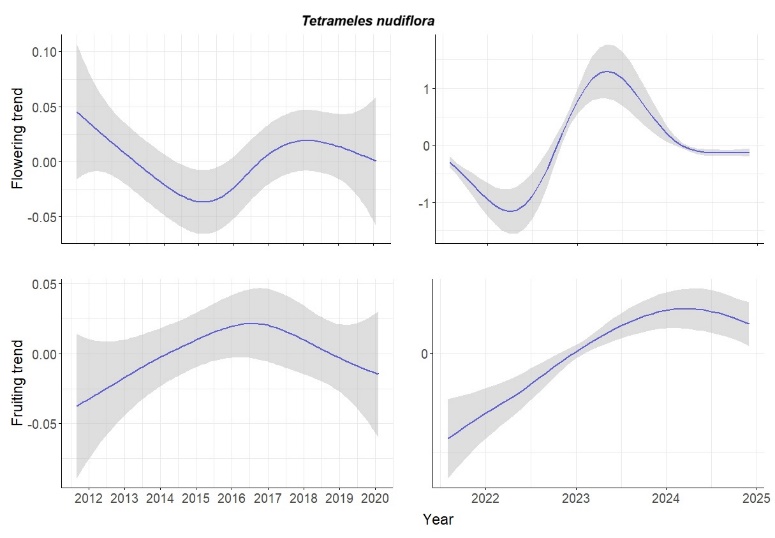

*Sterculia villosa	Tetrameles nudiflora*

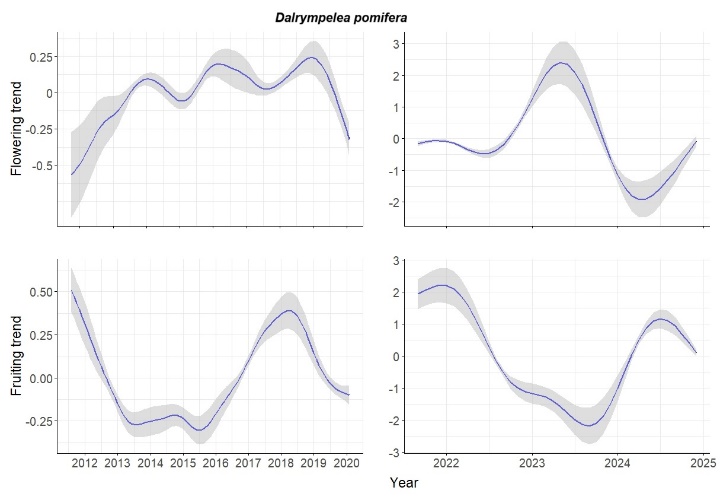

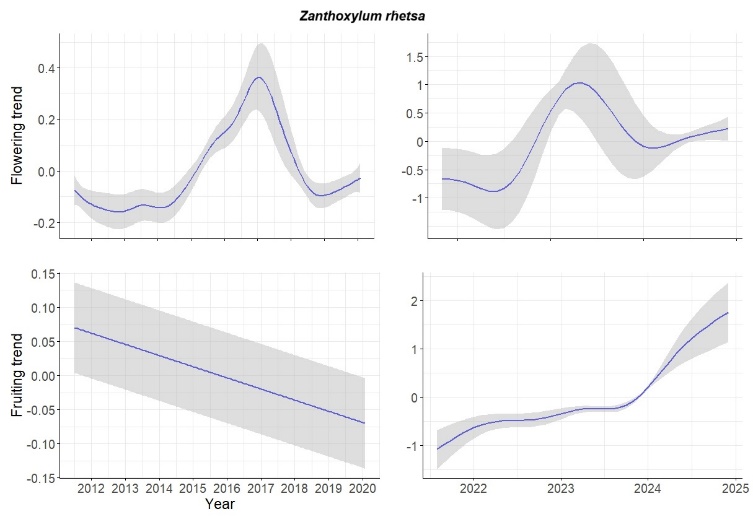

*Dalrympelea pomifera	Zanthoxylum rhetsa*

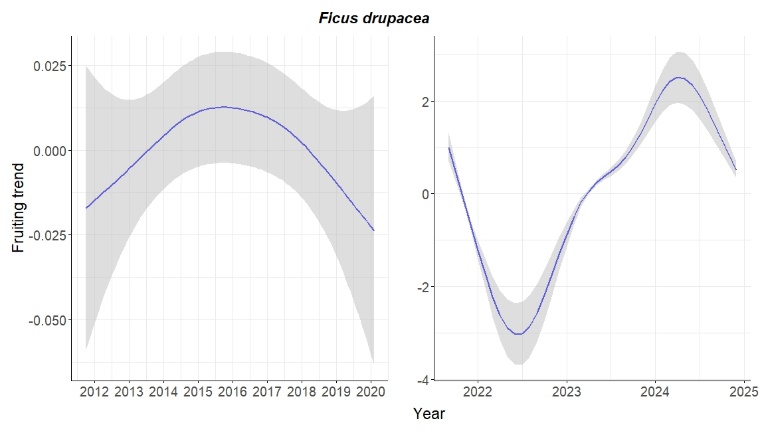

*Ficus drupacea*

Figure S1 : Smoothing splines of flowering and fruiting intensity of 36 tree species in Pakke Tiger Reserve, Arunachal Pradesh, India (for which at least 10 trees were monitored). Splines were fitted to trend components of phenological time series using Generalized Additive Models.

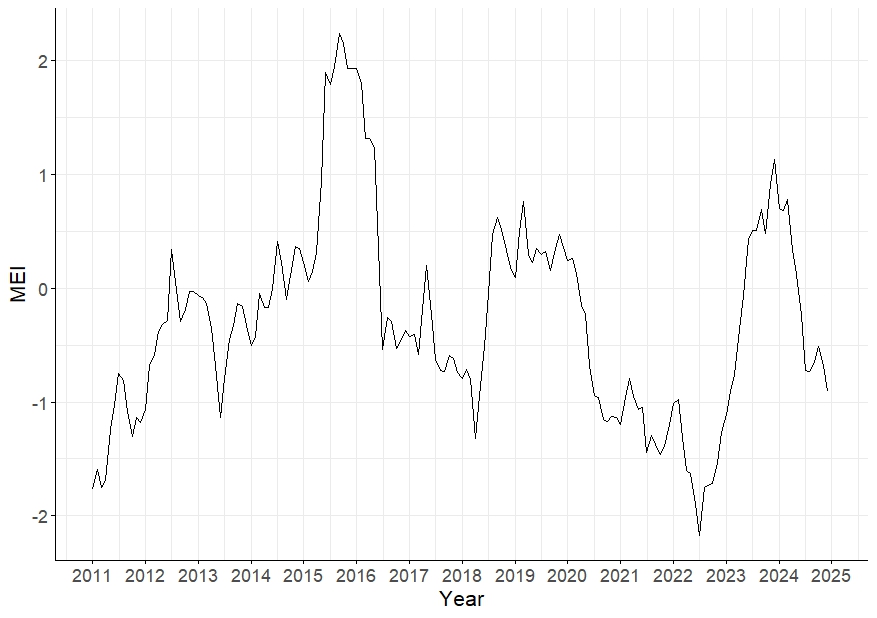

Figure S2: Trend in the MEI (Multivariate ENSO Index) in the period from January 2011 to December 2024.

Phenology-MEI CCFs

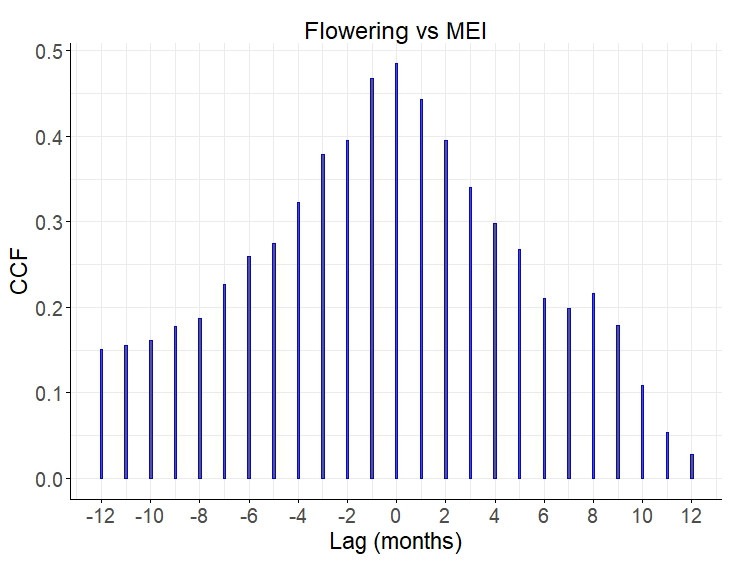

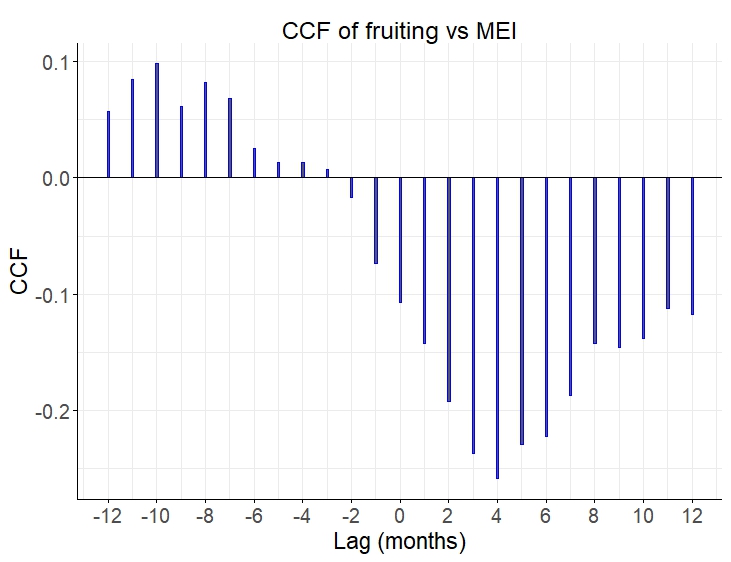

Climate-MEI CCFs

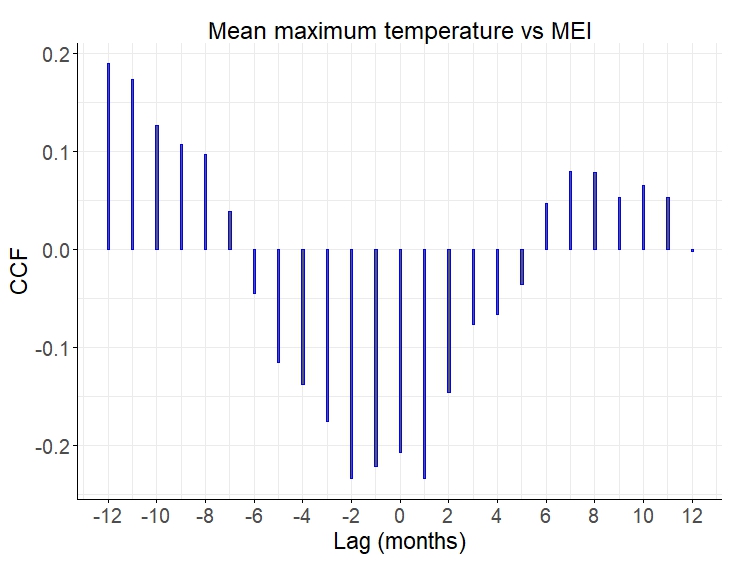

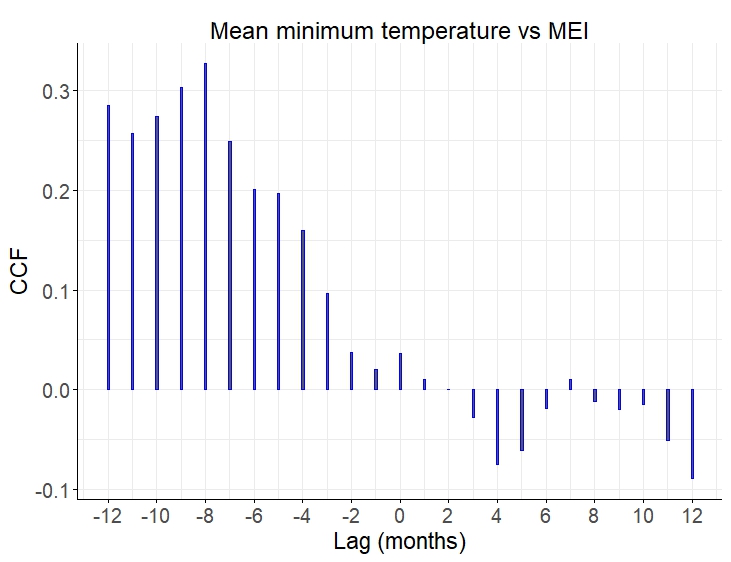

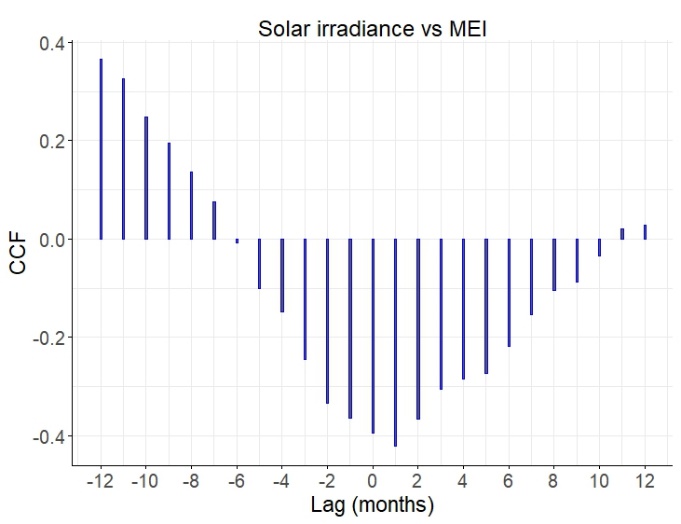

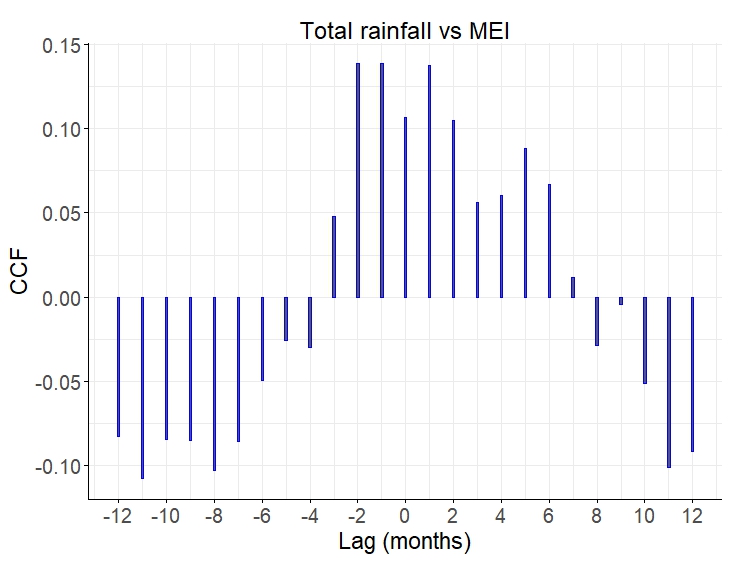

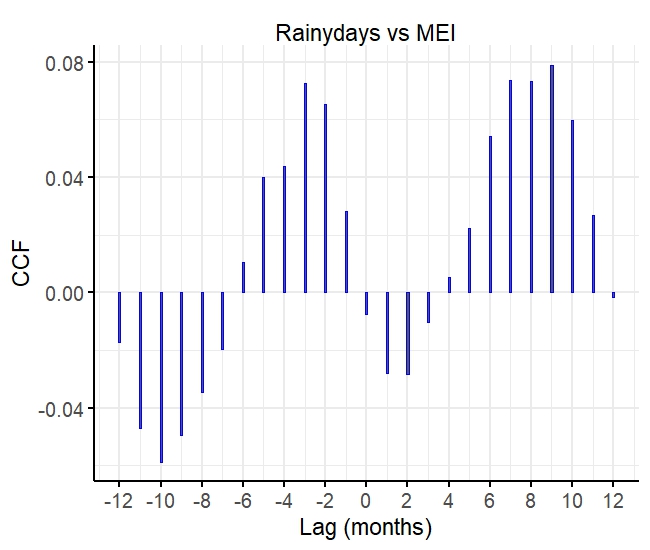

Climate-phenology CCFs

Flowering

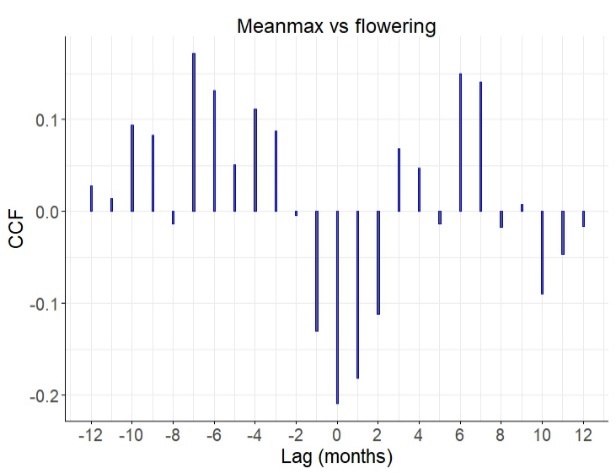

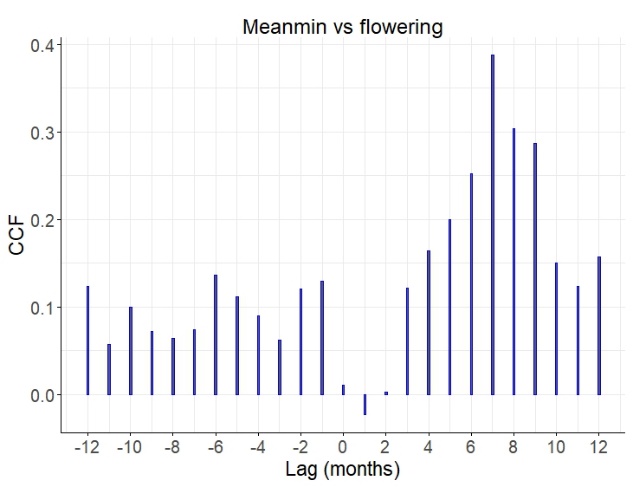

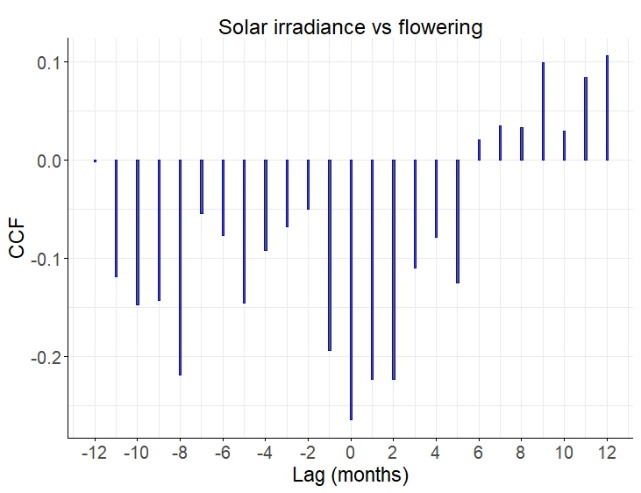

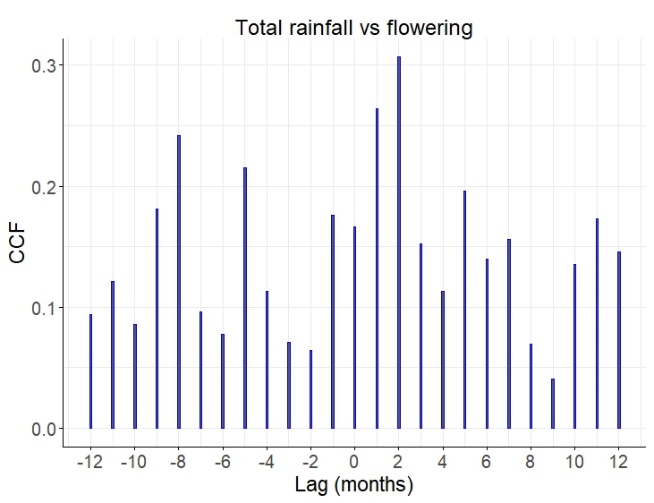

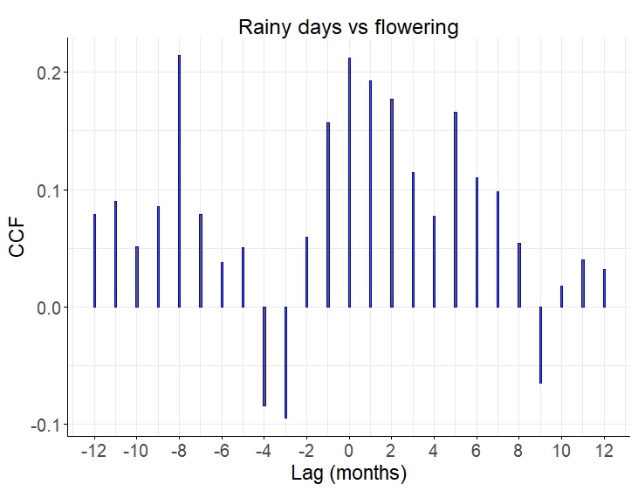

Fruiting

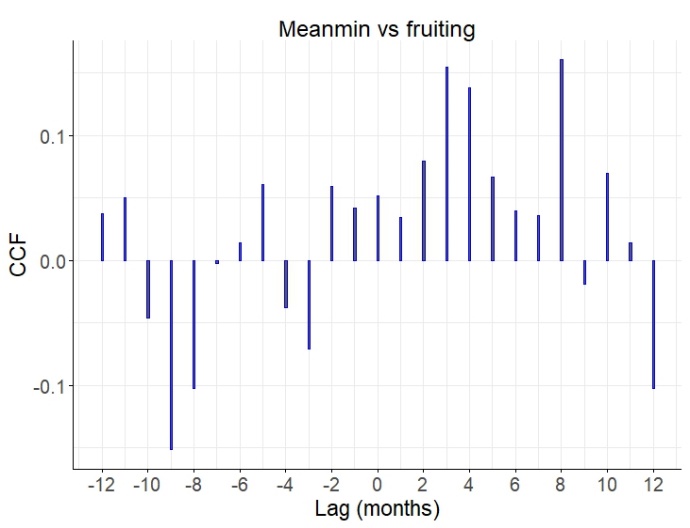

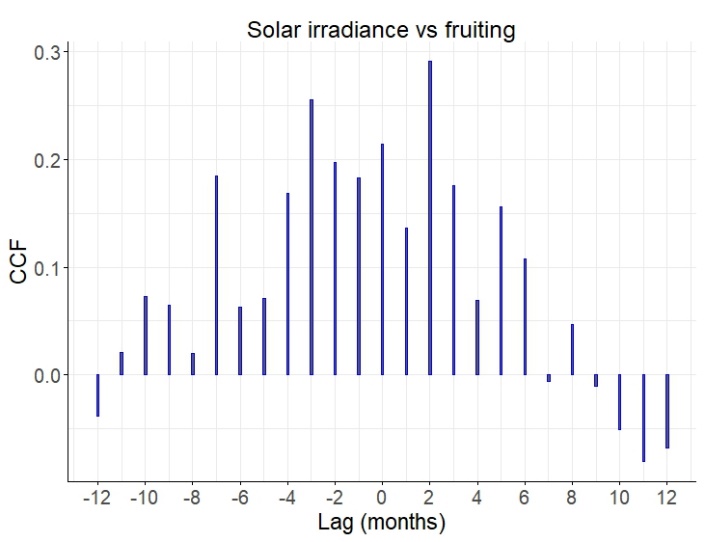

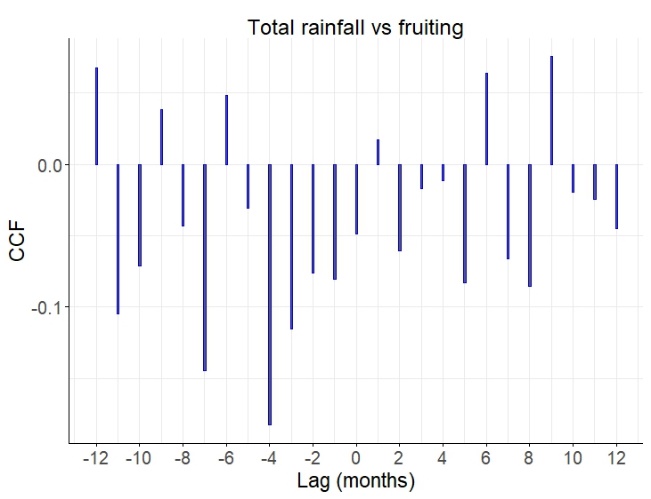

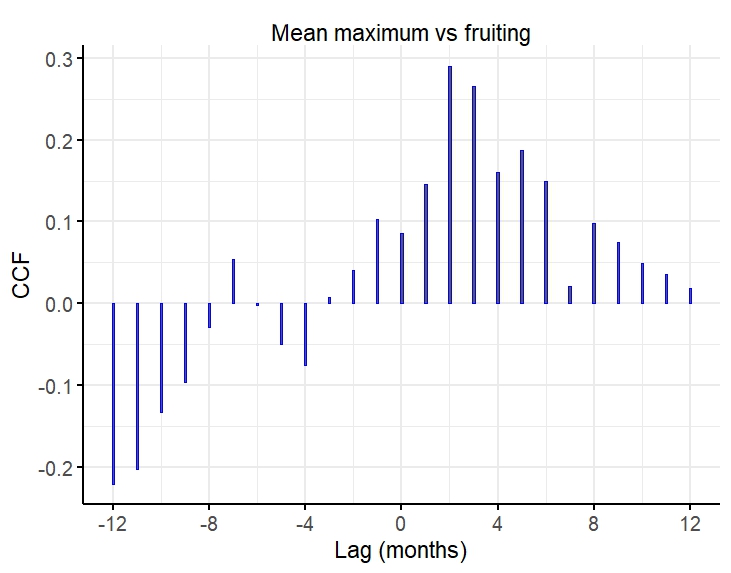

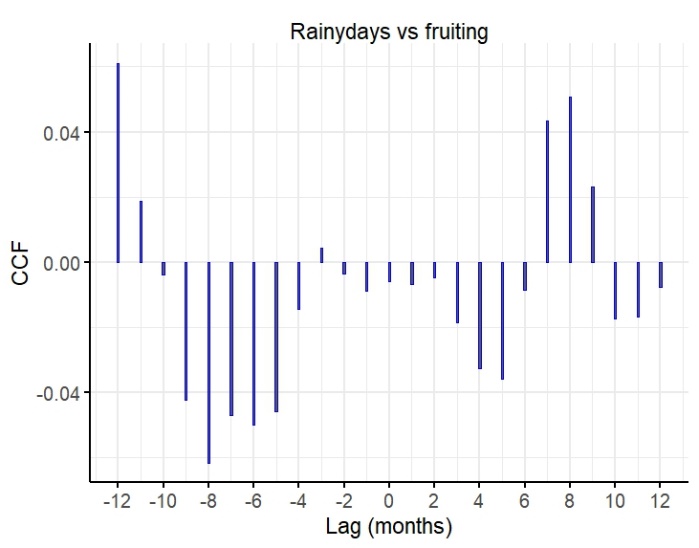

Fig. S3: Cross-correlation functions representing correlations between pairs of timeseries pertaining to MEI (Multivariate ENSO Index), temperature, precipitation and solar irradiance variables and phenological time series of flowering and fruiting. Cross-correlations represent lagged (in negative) effects and advanced (in positive) effects from 0-12 months.
